# Endogenously produced hydrogen cyanide serves as a novel mammalian gasotransmitter

**DOI:** 10.1101/2024.06.03.597146

**Authors:** Karim Zuhra, Maria Petrosino, Lucia Janickova, Kelly Ascenção, Thibaut Vignane, Jovan Petric, Moustafa Khalaf, Thilo M. Philipp, Stella Ravani, Abhishek Anand, Vanessa Martins, Sidneia Santos, Serkan Erdemir, Sait Malkondu, Barbara Sitek, Taha Kelestemur, Anna Kieronska-Rudek, Tomas Majtan, Luis Filgueira, Darko Maric, Stefan Chlopicki, David Hoogewijs, György Haskó, Andreas Papapetropoulos, Brian A. Logue, Gerry R. Boss, Milos R. Filipovic, Csaba Szabo

## Abstract

Small, gaseous molecules, known as gasotransmitters (NO, CO, H_2_S), are produced endogenously in mammalian cells and serve important biological roles. Hydrogen cyanide, traditionally considered a cytotoxic molecule in mammals, serves as an endogenous mediator in several plants and bacterial species. Here we show that low concentrations of cyanide are generated endogenously in mouse liver and human hepatocytes. Cyanide production is stimulated by glycine, occurs at the low pH of lysosomes and requires peroxidase activity. Cyanide, in turn, is detectable in several cellular compartments. Cyanide is also detectable basally in the blood of mice; its levels increase after treatment of the animals with glycine. Rhodanese activity regulates endogenous cyanide levels. Cyanide, when generated endogenously at an optimal level, exerts stimulatory effects on mitochondrial bioenergetics, cell metabolism and cell proliferation. Dysregulation of endogenous cyanide, either below or above optimal levels, impairs cellular bioenergetics. The regulatory effects of cyanide are in part mediated by posttranslational modification of cysteine residues via protein cyanylation; cyanylated protein residues can be detected basally, and increase after treatment with glycine. Controlled low-dose cyanide supplementation exhibits cytoprotective effects, as demonstrated in hypoxia and reoxygenation models *in vitro* and *in vivo*. However, pathologically elevated cyanide production, as demonstrated in nonketotic hyperglycinemia – an autosomal recessive disease of glycine metabolism – is deleterious to the cells.

---

Nitric oxide (NO), carbon monoxide (CO) and hydrogen sulfide (H_2_S) are small endogenous gaseous signaling molecules known as gasotransmitters**^1–7^**. Hydrogen cyanide is recognized as an endogenous regulator in bacteria and plants. However, in mammalian cells and tissues, it is generally regarded as a cytotoxic molecule**^8^**. In this study we investigated whether cyanide is produced in mammalian cells and tissues and, if so, whether it serves regulatory roles.*

## Cyanide is enzymatically produced in mammalian cells and tissues

To investigate cyanide formation in mammalian cells and tissues, we first used a cyanide-selective electrode**^9^**. This method is based on trapping gaseous cyanide via alkalization and the subsequent selective detection of CN^-^. Significant cyanide generation was detected from gently homogenized mouse tissues, with the liver producing the highest amounts (Fig. **1a**). We then tested whether amino acids could stimulate cyanide generation. Adding glycine to liver or spleen homogenates increased cyanide generation (Fig. **1a**, Extended Data Fig. 1a-e), while none of the other 19 proteinogenic amino acids exerted such response (Extended Data Fig. 1e). We confirmed the glycine-mediated stimulation of cyanide generation in liver homogenates using two additional methods which measure volatile HCN in the gaseous form: an LC-MS/MS method where HCN is trapped in a chamber containing naphthalene dialdehyde and taurine**^10^** (Fig. **1b**, Extended Data Fig. 1h,i), and a spectroscopic method, where HCN gas released from the tissue is captured by the cyanide scavenger monocyano-cobinamide (MCC), followed by a color change**^11^** (Fig. **1c**, Extended Data Fig. 1j-m). The specificity of the cyanide signal was confirmed by using the cyanide scavengers trihistidyl-cobinamide (THC) and dicobalt edetate (CoE)**^12,13^** (Fig. **1d**, Extended Data Fig. 1f).

**Figure 1.**
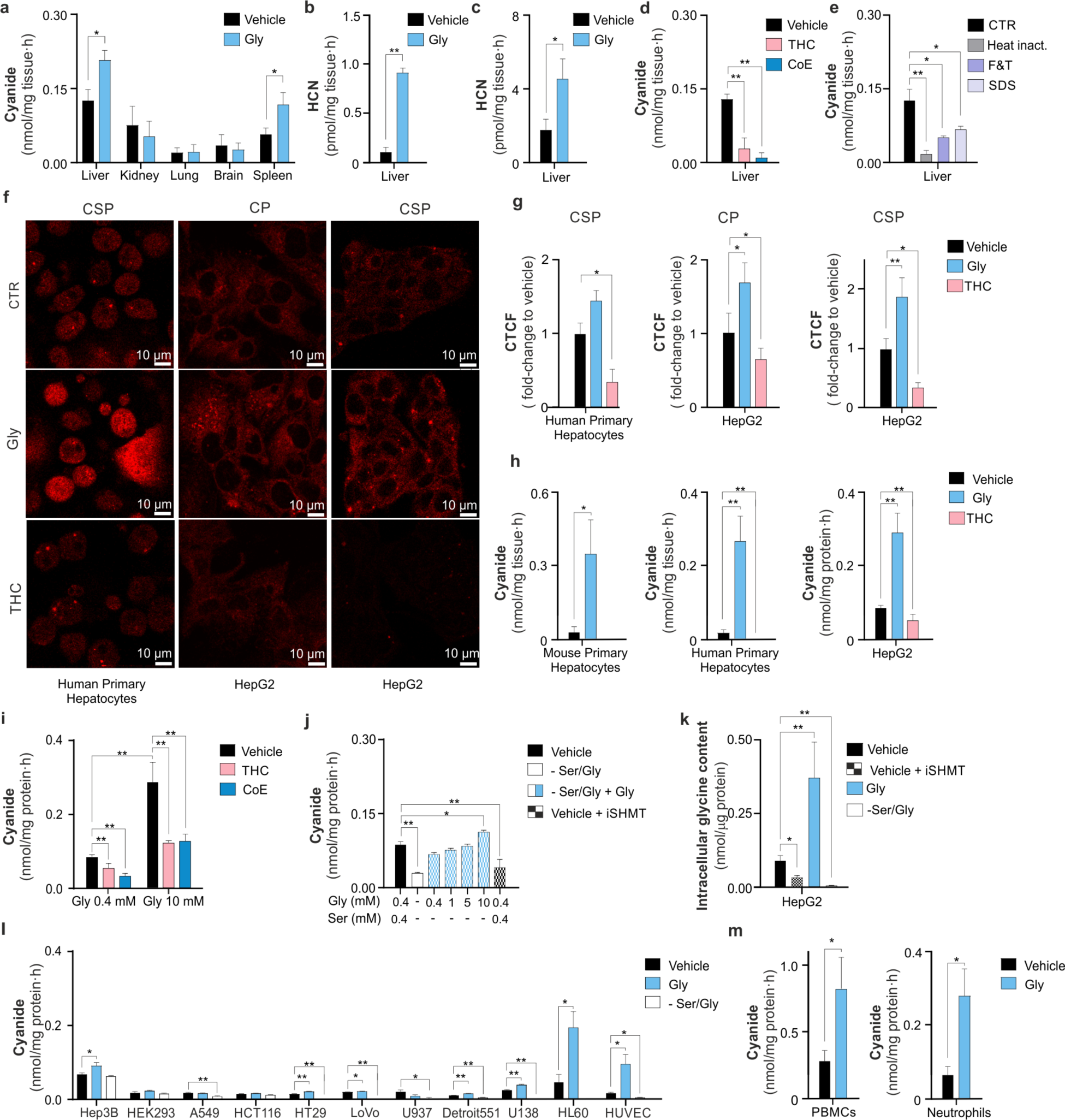
Cyanide is endogenously produced in mouse tissues and human cells. Cyanide generated by tissue homogenates was measured using **(a)** an electrochemical method (ECh) in which cyanide was trapped with NaOH, **(b)** an LC-MS/MS method where cyanide diffuses into a chamber and trapped as HCN and **(c)** a spectrophotometric method based on the reaction of HCN with a cyanide scavenger. Cyanide generation is shown in homogenization medium (Vehicle) or in medium supplemented with 10 mM glycine (Gly). **(d)** Treatment with HCN scavengers trihistidyl-cobinamide (THC) or dicobalt edetate (CoE) (10 µM), lowered the cyanide signal (ECh method) from mouse liver homogenates. **(e)** Heat-inactivation (Heat inact.), physical inactivation by multiple cycles of freezing and thawing (F&T) or SDS-induced protein denaturation (SDS) lowered the cyanide signal (ECh method) from mouse liver homogenates, compared to the control (CTR). **(f, g)** Corrected Total Cell Fluorescence (CTCF) values representing cyanide production in human primary hepatocytes and human hepatoma line (HepG2) detected with confocal microscopy using two different cyanide-sensitive fluoroprobes Chemosensor P (CP) and CSP, a spiropyrane derivative of cyanobiphenyl; the signal was detectable in control cells (CTR), increased in cells treated with glycine (Gly, 10 mM), and was reduced in cells treated with THC (10 µM). **(h)** Cyanide production in primary mouse and human hepatocytes and HepG2 cells (ECh method); the signal was increased in cells treated with glycine (Gly, 10 mM), and was reduced in cells treated with THC (10 µM). **(i)** Effect of THC or CoE (10 µM) on the cyanide signal in HepG2 cells (ECh method). **(j)** Cyanide production in HepG2 cells grown for 24 h in normal medium ± 100 µM of a SHMT inhibitor or in Ser/Gly-free medium supplemented with increasing concentrations of glycine (400 µM – 10 mM) (ECh method). **(k)** Glycine levels in HepG2 cells under baseline conditions, after pharmacological inhibition of SHMT (iSHMT), after addition of glycine (10 mM) to the culture medium or after omission of serine/glycine (-Ser/Gly) from the medium for 24h. **(l)** Cyanide production in a panel of mammalian cell lines in normal medium, in medium supplemented with 10 mM glycine (Gly) or when maintained in Ser/Gly-free medium (-Ser/Gly) for 24 hours. **(m)** Cyanide production in human peripheral blood mononuclear cells (PBMCs) and human neutrophils under basal conditions and after incubation with glycine (Gly) for 4 hours. Data show mean±SEM of at least n=5 independent experiments. *p<0.05 or **p<0.01 indicate significant differences.

Having shown that glycine serves as a stimulator of mammalian cyanide generation, we aimed to determine if this process is enzymatically regulated. We found that cyanide generation was abolished by several different methods that denature of the proteins in the tissue homogenate, suggesting the involvement of an enzymatic process (Fig. **1e**).

Human and mouse primary hepatocytes and the human hepatoma cell line (HepG2) also produced cyanide when assayed in standard culture medium containing 400 µM glycine (Fig. **1f-k**). The cyanide signal increased by glycine supplementation and reduced by the cyanide scavenger THC, as assessed by confocal microscopy using two structurally different fluorescent cyanide probes**^14,15^** (Fig. **1f,g**) and also quantified by the electrochemical method (Fig. **1h,i**, Extended Data Fig. 2a,b). Importantly, cyanide generation was suppressed in cells grown in serine/glycine-free (- Ser/Gly) medium, and re-addition of glycine restored cyanide generation. (Fig. **1j**). The clinically used drug iclepertin, which inhibits the glycine transporter GlyT-1 (SLC6A9) on the cell membrane also inhibited cyanide generation (Extended Data Fig. 2c). Cyanide production was also attenuated by treatment of the cells with a pharmacological inhibitor of serine hydroxymethyltransferase (SHMT)**^16^**, the enzyme which interconverts glycine and serine (Fig. **1j**) and a similar inhibitory effect was also observed with another SHMT inhibitor, lometrexol (Extended Data Fig. 2d). Measurement of intracellular glycine concentrations (Fig. **1k**) confirmed the modulation of intracellular glycine levels by the above interventions.

Collectively, these data indicate that glycine is an endogenous stimulator of cyanide generation. Cyanide generation was detectable from various cultured human cell lines including cells of lung and colonic epithelial and myeloid lineage; the cells responded to glycine supplementation or deprivation with increased or decreased cyanide production, respectively (Fig. **1l**). Moreover, significant basal cyanide production was detectable in human peripheral blood mononuclear cells (PBMCs) and neutrophils, and cyanide generation was markedly further stimulated after incubation with glycine (Fig. **1m**). From the parenchymal cells investigated, cells of hepatic origin, including primary hepatocytes, HepG2, and Hep3B, exhibited the highest levels of cyanide production (Fig. **1i-l**). Therefore, we selected the human hepatoma line HepG2 for subsequent studies.

## Cyanide production occurs in lysosomes

Confocal live cell imaging revealed that cyanide is present throughout the cell, as expected for a gaseous molecule with free diffusion (Fig. **1f**). Thus, cyanide was detectable in the cytosol – as evidenced by partial co-localization with calcein AM, and also in mitochondria – as evidenced by partial co-localization with the mitochondrial marker MitoTracker (Extended Data Fig. 3a,b). However, the strongest cyanide signal was detected in the lysosomes, as evidenced by co-localization with Lysotracker (Fig. **2a**).

**Figure 2.**
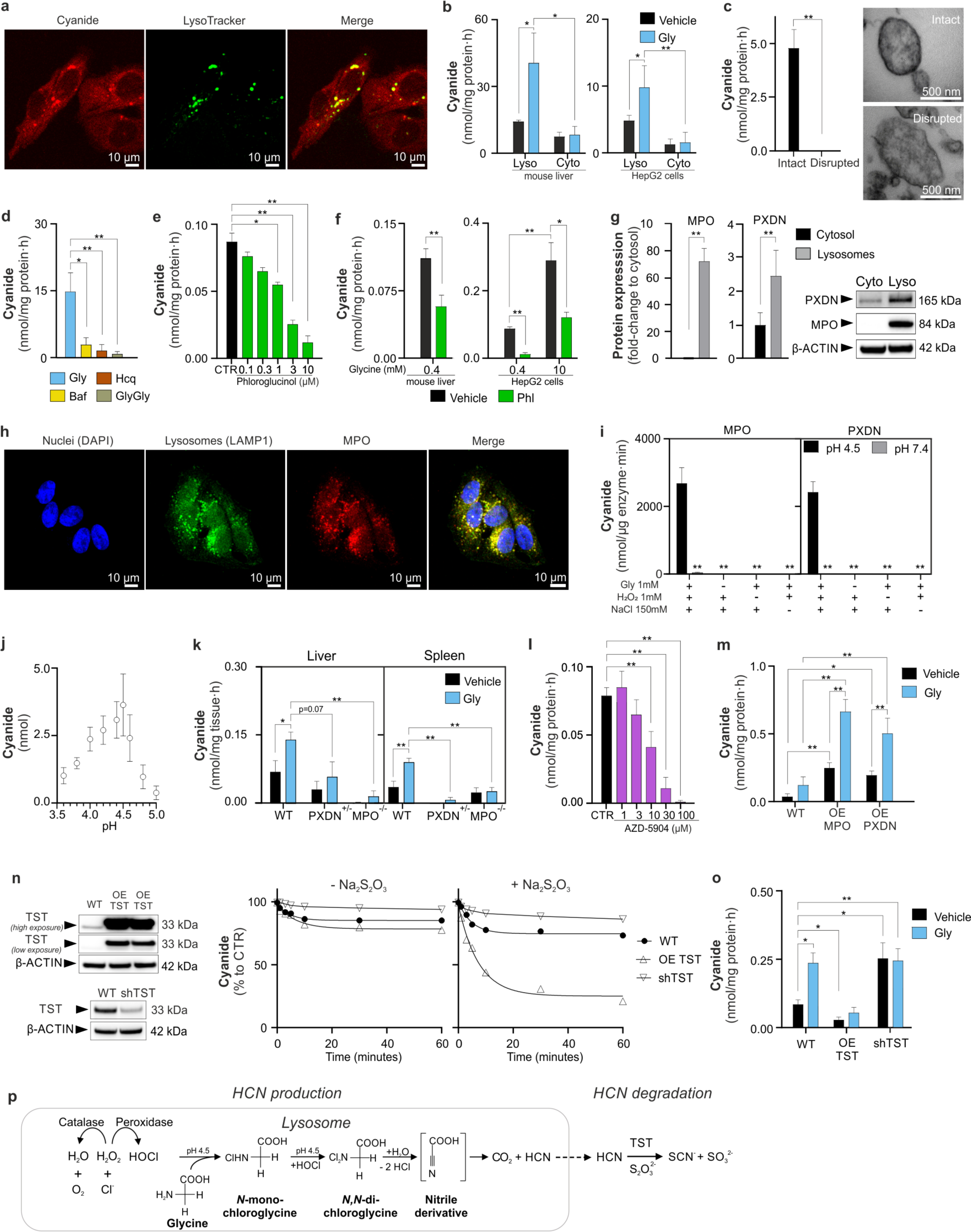
Cyanide is enzymatically generated by lysosomal peroxidases. **(a)** The cyanide signal in HepG2 cells partially colocalizes with lysosomes (confocal microscopy, using Chemosensor P). **(b)** Cyanide generation in lysosomal and cytosolic fractions (Lyso, Cyto, respectively) obtained from mouse liver or HepG2 cells <u>±</u> glycine (Gly, 10 mM) or **(c)** from isolated intact vs. disrupted lysosomes (ECh method). Lysosome disruption was confirmed by electron microscopy. **(d)** Cyanide generation (ECh method) in isolated lysosomes after treatment with bafilomycin (Baf, 1 µM), hydroxychloroquine (Hcq, 30 µM) or glycine-glycine dipeptide (Gly-Gly, 150 mM). **(e)** Cyanide detection in HepG2 cells or **(f)** mouse liver homogenates treated with various concentrations (0.1-10 µM) of phloroglucinol (Phl) for isolated lysosomes or with a single concentration (10 µM) for mouse liver homogenates and HepG2 cells (ECh method). **(g)** Myeloperoxidase (MPO) and peroxidasin (PXDN) expression in lysosomal (L) and cytosolic (C) fractions of HepG2 cells (Western blotting). **(h)** MPO localizes to lysosomes (confocal microscopy). Nuclei were visualized with 4′,6-diamidino-2-phenylindole (DAPI), lysosomes with a lysosomal associated membrane protein 1 (LAMP-1 antibody), and MPO with an MPO antibody **(i)** MPO or PXDN catalyze cyanide generation at pH 4.5 (biochemical assay, enzyme was incubated with various combinations of glycine (Gly, 1 mM), hydrogen peroxide (H_2_O_2_, 1 mM) and NaCl (150 mM) (ECh method). **(j)** Equimolar concentrations of HOCl and glycine generate cyanide (pH optimum: 4.5) (biochemical assay, ECh method). **(k)** Cyanide production in liver and spleen homogenates of wild-type (WT), PXND^+/-^ and MPO^-/-^ mice under baseline conditions and after the addition of glycine (Gly, 10 mM) (ECh method). **(l)** Detection of cyanide in HepG2 cells treated with the MPO inhibitor 1,2,3,9-tetrahydro-3-[[(2R)-tetrahydro-2-furanyl]methyl]-2-thioxo-6H-purin-6-one (AZD-5904, 1-100 µM) and **(m)** in HEK293T cells overexpressing (OE) MPO or PXDN (ECh method). **(n)** Overexpression of human rhodanese (OE TST) or its downregulation (shTST) in HepG2 cells accelerates or reduces, respectively, the cellular degradation of exogenously administered KCN. **(o)** Overexpression of human rhodanese (OE TST) or its downregulation (shTST) in HepG2 cells results in the reduction or accumulation, respectively of endogenous cyanide in HepG2 cells (ECh method). Data are presented as mean±SEM of at least n=5 independent experiments. *p<0.05 or **p<0.01 indicate significant differences. **(p)** Proposed scheme of lysosomal cyanide generation. In lysosomes, at pH 4.5, glycine undergoes a two-step chlorination reaction in presence of peroxidases-derived HOCl. The subsequent hydrolysis of *N,N*-dichloroglycine leads to the formation of an unstable nitrile derivative intermediate which spontaneously decomposes to carbon dioxide (CO_2_) and hydrogen cyanide (HCN). HCN, in turn, is converted to SCN^-^ and CO_2_ via rhodanese/TST in the extralysosomal compartments.

Because the cyanide signal was most prominent in the lysosomes, we next compared the cyanide-generating capacity of isolated lysosomal vs. cytosolic fractions obtained from liver tissue and HepG2 cells. Higher baseline cyanide concentrations were detected in the lysosomal fraction than in the cytosolic fraction; adding glycine selectively increased cyanide generation in the lysosomal – but not the cytosolic – fraction (Fig. **2b**, Extended Data Fig. 4a-c). Electron microscopy confirmed that our standard homogenization protocol preserved the structural integrity of subcellular organelles, including lysosomes (Fig. **2c**). In contrast, a more drastic homogenization protocol – which disrupted lysosomes – abrogated the ability of glycine to stimulate lysosomal cyanide generation (Fig. **2c**), suggesting that the integrity of the lysosomes is essential for mammalian cyanide generation.

We confirmed the importance of functional lysosomes for cyanide generation by demonstrating that the lysosomal proton pump inhibitor bafilomycin (Baf)**^17^**, the lysosomal alkalinizer^b^ hydroxychloroquine (Hcq)**^18^** and Gly-Gly dipeptide, which inhibits the lysosomal glycine transporter LYAAT-1**^19^**: all of these interventions suppressed cyanide generation in isolated lysosomes (Fig. **2d**); bafilomycin and hydroxychloroquine also suppressed cyanide generation in HepG2 cells (Extended Data Fig. 4d-f).

## Cyanide production requires peroxidase activity

Enzymatic conversion of glycine to cyanide would require oxidation of glycine’s amino group to a nitril group, something that hydrolyzing enzymes, the main constituents of lysosomes, cannot do. However, peroxidases, which may generate strong oxidants, such as hypochlorite (HOCl), may catalyze glycine’s oxidation. We therefore tested the effect of the broad-spectrum peroxidase inhibitor phloroglucinol**^20^** (Phl) on cyanide levels in liver homogenates and HepG2 cells and found that it concentration-dependently inhibits cyanide production (Fig. **2e,f**, Extended Data Fig. 4e). As in most cells**^21^**, multiple peroxidases were detected in HepG2 cells, with myeloperoxidase (MPO) and peroxidasin (PXDN) exhibiting preferential lysosomal localization (Fig. **2g**, Extended Data Fig. 4g). Confocal microscopy confirmed MPO localization to the lysosomes, but not to the endoplasmic reticulum or mitochondria (Fig. **2h**, Extended Data Fig. 3c,d).

Because mammalian cyanide production (i) requires glycine, (ii) occurs in the acidic pH of lysosomes and (iii) is peroxidase dependent, we hypothesized that cyanide generation is dependent on the peroxidase product hypochlorous acid (HOCl), which is predominantly produced in lysosomes**^22^**. Indeed, we found that HOCl mainly localized to the lysosomes, and to a lesser extent to the cytosol of HepG2 cells (Extended Data Fig. 3d). We, therefore, conducted *in vitro* biochemical experiments, where we incubated MPO or PXDN enzyme with glycine, hydrogen peroxide (H_2_O_2_) and chloride (Cl^-^), and observed cyanide production with a pH optimum of 4.5 (Fig. **2i**). Additionally, when glycine was incubated with HOCl in the absence of any enzyme, cyanide was also produced with the same pH optimum of 4.5 (Fig. **2j**). Other proteinogenic amino acids did not generate significant amounts of cyanide under the same conditions (Extended Data Fig. 5a).

Liver and spleen homogenates obtained from MPO^-/-^ or PXDN^+/-^ mice generated less cyanide than homogenates from wild-type animals (Fig. **2k**, Extended Data Fig. 5b,c). The MPO inhibitor AZD-5904**^23^** also inhibited cyanide generation (Fig. **2l**). Overexpression of MPO or PXDN in HEK293 cells markedly increased (Fig. **2m**), while overexpression of catalase reduced cyanide generation (Extended Data Fig. 5d). Together, these data indicate that peroxidase-catalyzed glycine oxidation in lysosomes is the predominant mechanism of endogenous cyanide generation in mammalian cells (Extended Data Fig. 5g).

Thiosulfate sulfur transferase (TST, also known as rhodanese) is considered the main cyanide detoxifying enzyme in eukaryotes**^24^**. Overexpression of this enzyme in HepG2 cells increased the ability of the cells to decompose exogenously added cyanide (applied as the salt form, KCN to the cells) (Fig. **2n**) and decreased endogenous cyanide concentrations in HepG2 cells (Fig. **2o**). Similar effects were observed when the bacterial cyanide degradation enzyme cyanide dihydratase (CynD)**^25^** was overexpressed (Extended Data Fig. 5e,f). Conversely, knockdown of the TST gene (shTST) resulted in reduced KCN degradation rates (Fig. **2n**) and increased the accumulation of endogenous cyanide (Fig. **2o**).

Based on all these results, combined with prior biochemical findings**^26^**, we propose the following model of mammalian cyanide generation (Fig. **2p**): peroxidases – MPO, PXDN and possibly others – produce HOCl, using H_2_O_2_ and chloride as substrates; HOCl, in the acidic milieu of the lysosome, reacts with glycine, yielding N-monochloroglycine, which undergoes acid-catalyzed conversion to N,N-dichloroglycine. The latter molecule decomposes to the corresponding nitrile, cyanocarboxylic acid (CN-COOH), releasing hydrogen cyanide and CO_2_. Due to its gaseous properties, cyanide reaches various intracellular components, and may also diffuse out of the cell and act as a paracrine mediator, thereby serving as a mammalian gasotransmitter (Extended Data Fig. 5g).

## S-cyanylation is induced by endogenously generated cyanide in mammalian cells

Gasotransmitters regulate various cellular functions via posttranslational modifications (PTMs) of cysteine, such as cysteine S-nitrosylation by NO or cysteine persulfidation by H_2_S**^1–7^**. Cyanylation (RSCN), the addition of a CN group to the sulfur atom in cysteine residues, has been recently described in plants**^27^**. This posttranslational modification has been suggested to affect protein function and has been proposed to influence various signaling pathways and cellular responses**^27^**.

Therefore, we evaluated proteins from mouse liver and HepG2 cells for evidence of cyanylation. In the absence of a chemoselective method to assess cyanylation, we analyzed proteome data for loss of a hydrogen atom and addition of a CN group (m/z +24.995249 Da). We lysed tissues or cells with iodoacetamide to block all available thiols and found cyanylation of 249 cysteines on 214 proteins in mouse liver under baseline conditions. Incubation of liver lysates with 10 mM glycine increased cyanylation up to 10-fold on 71 cysteine sites (Fig. **3a**, Extended Data Table 1). Since the total number of cysteine PTMs are unknown, and that some of them are not reducible, it is not possible to estimate the cysteine site occupancy by cyanylation. However, we did quantify the change of reduced thiols (which should still represent the most abundant form of cysteine) in those glycine-treated liver extracts and observed that where cyanylation increased, free thiols decreased (Fig. **3a,b**, Extended Data Table 1). These data indicate that a significant portion of the total proteome is modified by cyanylation. The glycine-induced increase in cyanylated proteins in liver was greatest in the lysosomes (Fig. **3c**), consistently with lysosomes being the source of cyanide in mammalian cells. Cyanylation also affected multiple proteins involved in glycolysis, apoptosis, stress response and glutathione metabolism (Fig. **3c,d**, Extended Data Table 1). To confirm that cyanylation originated from C and N in glycine, we also treated liver lysates with ^13^C,^15^N-labelled glycine. In these experiments we detected 354 “heavy” cyanylated peptides on 201 proteins (Fig. **3e**, Extended Data Table 2).

**Figure 3.**
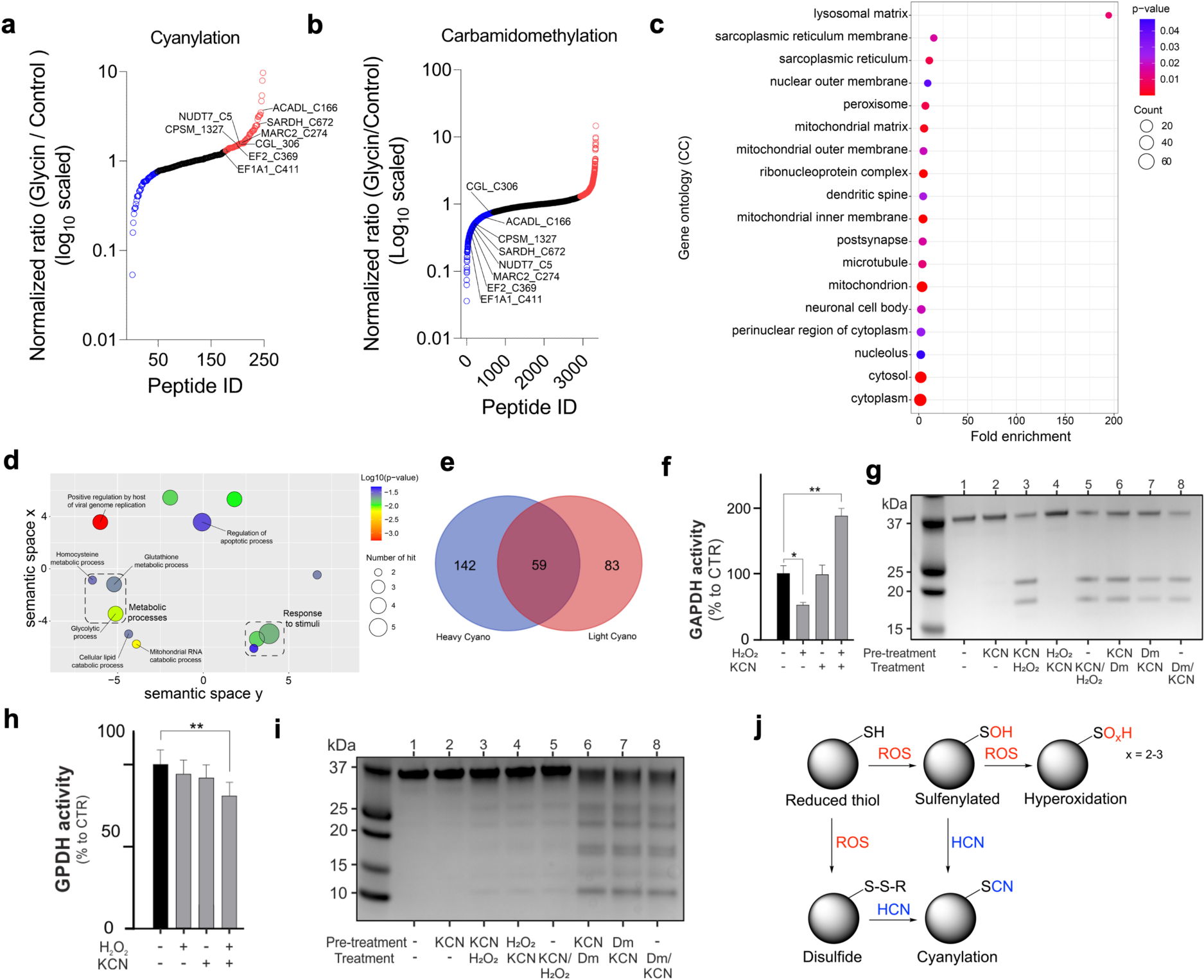
Cyanide induces posttranslational protein modifications. **(a)** The proteome-wide and site-specific changes in *S*-cyanylation in mouse liver tissue lysates treated with 10 mM glycine (Gly, n = 5). **(b)** The proteome-wide and site-specific changes in carbamidomethylation (reduced thiol labelled with iodoacetamide) in mouse liver tissue lysates treated with 10 mM Gly (n = 5). **(c)** GO term enrichment analysis (cellular localization) of proteins whose S-cyanylation increases in liver samples treated with 10 mM Gly. **(d)** GO term enrichment (biological process) of the proteins whose S-cyanylation is significantly increased in liver tissue treated with 10 mM Gly. Employing DAVID for enrichment, the outcomes were visualized through REVIGO. Significant GO terms passed the Benjamini adjusted p-value threshold of 0.01. Circle dimensions denote the protein count within specific GO terms, while color gradients communicate the degree of significance. **(e)** Venn diagram comparing the proteins found to contain ^13^C^15^N (heavy cyano) cyanylation with the endogenously cyanylated proteins (light cyano) in liver tissue lysates treated with ^13^C^15^N-labelled Gly (n = 5). **(f)** Enzymatic activity of glyceraldehyde 3-phosphate dehydrogenase (GAPDH) pre-incubated with H_2_O_2_ (10 µM), KCN (10 µM) or H_2_O_2_/KCN. Data are presented as mean ± SEM of at least n = 5 independent experiments. *p<0.05 or **p<0.01 indicate significant differences. **(g)** Detection of high pH-induced peptide bond cleavage at cyanylation sites of GAPDH that was treated different combination of KCN, H_2_O_2_ or diamide (Dm) (SDS-PAGE analysis). **(h)** Enzymatic activity of glycerol-3-phosphate dehydrogenase (GPDH) pre-incubated with H_2_O_2_ (10 µM), KCN (10 µM) or H_2_O_2_/KCN. Data are presented as mean ± SEM of at least n = 3 independent experiments. **p<0.01 indicate significant differences. **(i)** Detection of high pH-induced peptide bond cleavage at cyanylation sites of GPDH that was treated different combination of KCN, H_2_O_2_ or diamide (Dm) (SDS-PAGE analysis). **(j)** Proposed scheme of protein S-cyanylation. Upon reaction with reactive oxygen species (ROS) thiols (RSH) become oxidized to either sulfenic acid (RSOH) or to disulfides (RSSR) which could be both intra and intermolecular disulfides. Both, ROSH and RSSR could react with HCN leading to protein cyanylation (RSCN). When SH groups are hyperoxidized, they are no longer reactive to cyanide.

Cyanylation also increased in HepG2 cells supplemented with glycine (Extended Data Fig. 6a,b, Extended Data Table 3). Conversely, S-cyanylation of ∼60 sites decreased in HepG2 cells grown in Gly/Ser free medium, affecting proteins involved in telomere maintenance, response to stress, and regulation of protein ubiquitination (Extended Data Fig. 6c,d, Extended Data Table 3). Furthermore, we also observed increased protein cyanylation in HepG2 cells treated with exogenously administered cyanide (10 nM or 1 µM KCN) and with several cyanogenic molecules – amygdalin, linamarin or mandelonitrile (Extended Data Fig. 6e,f, Extended Data Table 4).

To better understand the mechanism of cyanylation and its potential outcome on enzyme activity we used glyceraldehyde-3-phosphate dehydrogenase (GAPDH) and glycerol-3-phosphate dehydrogenase (GPDH) as two model proteins. Incubation of GAPDH with H_2_O_2_ inhibited its catalytic activity, while inhibition with KCN had no effect. However, incubating the enzyme with equimolar amounts of H_2_O_2_ and KCN yielded almost a two-fold increase in enzyme activity (Fig. **3f**). Based on gel electrophoresis and MS experiments, we found evidence for S-cyanylation of cysteine residues based on characteristic peptide bond cleavage at cyanylation site upon alkalization (Fig. **3g**, Extended Data Fig. 7a-c). Contrary to GAPDH, where cyanylation increased enzymatic activity, cyanylation had an inhibitory effect on GPDH (Fig. **3h,i**), which was associated with the cyanylation of multiple residues (Extended Data Fig. 7d). Based on the known reactivity profile of cyanide with thiol groups**^27–29^**, we propose that cyanide reaction with cysteine residues requires that the SH group is first oxidized to either sulfenic acid (-SOH) or to a disulfide (S-S or S-SR) and then cyanide reacts to yield the SCN product (Fig. **3j**). GPDH and GAPDH were used as model proteins to study the functional effects of cyanylation, but it is important to point out that in our proteome data we have also identified both GAPDH and GPDH as proteins whose cyanylation increases ∼50% (C245) and ∼75% (C546), respectively, upon treatment with glycine (Extended Data Table 1,2).

Taken together, the above data demonstrate that (i) mammalian proteins are endogenously cyanylated, (ii) cyanylation originates from glycine as a source of cyanide and could occupy a substantial portion of protein’s thiol pool, and (iii) cyanide can remodel the intracellular landscape of cysteine posttranslational modifications, with a functional outcome (e.g. enzyme activation or inactivation); cyanide may also serve as a redox switch from one post-translational modification (such as SH oxidation or glutathionylation) to another (cyanylation).

## Physiological cyanide generation supports cellular bioenergetics and proliferation

Based on the known roles of NO, CO, and H_2_S in mammalian cells**^1–7^** and on the recently demonstrated stimulatory bioenergetic effects of low concentrations of exogenously administered KCN**^30^**, we tested the role of endogenously-generated cyanide on bioenergetics and cell proliferation. Treatment of HepG2 cells with glycine increased mitochondrial electron transport and ATP generation and cyanide scavengers abrogated glycine’s stimulatory effect (Fig. **4a**, Extended Data Fig. 8a). Serine/glycine deprivation also exerted inhibitory effects on various bioenergetic parameters (Fig. **4a**, Extended Data Fig. 8a,b).

**Figure 4.**
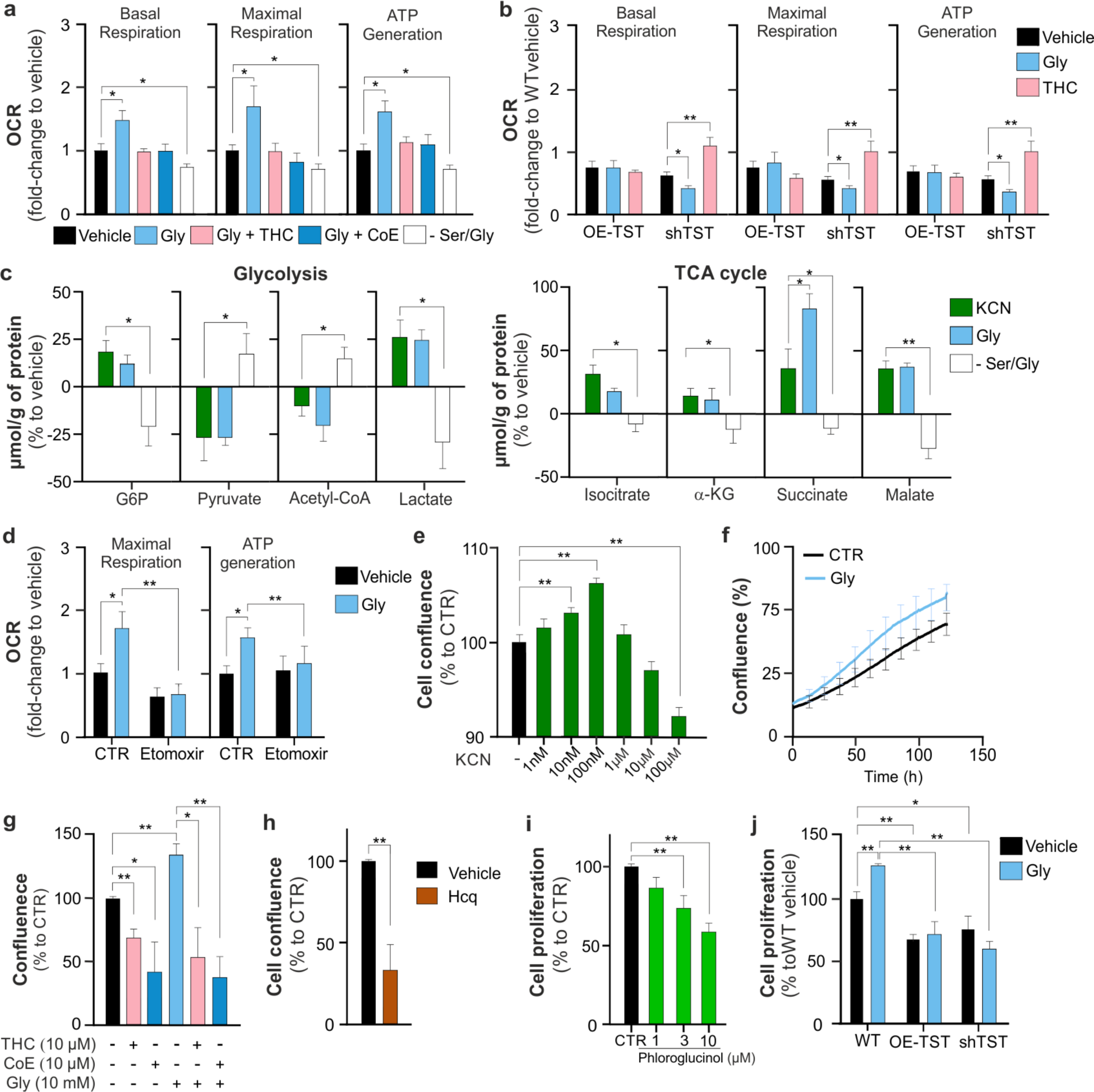
Endogenous cyanide generation supports cellular bioenergetics and proliferation. **(a)** Bioenergetic profile of HepG2 cells treated with glycine (Gly, 10 mM, 24 h), cyanide scavengers (10 µM, 3 h), or Ser/Gly deprivation. Oxygen consumption rates (OCR) are shown. **(b)** Bioenergetic profile of wild-type *vs.* TST-overexpressing (OE-TST) *vs.* TST-knocked-down (shTST) HepG2 cells ± glycine (10 mM, 24 h). OCR are shown. **(c)** Metabolomic analysis of HepG2 cells subjected to exogenous KCN (10 nM), glycine (10 mM) or serine/glycine deprivation for 24 h. G6P: glucose-6-phosphate. **(d)** Free fatty acid (FFA) oxidation analysis of HepG2 cells ± glycine (10 mM, 24 h). OCR are shown. Proliferation of HepG2 cells **(e)** in the presence of various concentrations of KCN (1 nM - 100 µM) **(f)** supplemented with glycine (10 mM) over 120 hours, measured at the IncuCyte system **(g)** in the presence of 0.4 mM (control level) or 10 mM glycine in the presence or absence of trihistidyl-cobinamide (THC,10 µM) or dicobalt edetate (CoE, 10 µM), **(h)** hydroxychloroquine (Hcq, 10 µM), or **(i)** phloroglucinol (1-10 µM) recorded at 24 hours. **(j)** Proliferation of wild-type *vs.* TST-overexpressing (OE-TST) *vs.* TST-knocked-down (shTST) HepG2 cells in the absence or presence of 10 mM Gly supplementation, recorded at 24 hours. Data are presented as mean±SEM of at least n=5 independent experiments. *p<0.05 or **p<0.01 indicate significant differences.

The effect of glycine on cellular bioenergetics was also attenuated in cells overexpressing either TST or CynD (Fig. **4a,b**, Extended Data Fig. 8c,d). In TST overexpressor cells, basal bioenergetic parameters were approximately 30% lower than in wild-type control cells (Fig. **4b**). Importantly, basal cellular bioenergetics was also diminished – by approximately 40% – in cells where TST was knocked down (shTST) (Fig. **4b**). In the shTST cells the cyanide scavenger THC improved bioenergetic function (Fig. **4b**), while in TST overexpressor cells addition of glycine or scavenging of cyanide did not affect bioenergetic parameters (Fig. **4b**). The above data suggest that – similar to the bell-shaped concentration-responses associated with NO, CO, and H_2_S**^1–7^** – endogenously-produced cyanide supports cellular bioenergetics with a concentration optimum: cellular bioenergetics is impaired both when endogenous cyanide levels are decreased, and when these levels are increased above beyond the optimal levels. Indeed, the inhibition of cytochrome C oxidase and the consequent suppression of mitochondrial function in response to high concentrations of cyanide are well-established facts in the toxicological literature**^8,34^**.

Metabolomic analysis demonstrated that glycine – similar to a low concentration of KCN – stimulates glycolysis and activates the Krebs cycle, while serine/glycine deprivation exerts inhibitory effects (Fig. **4c**, Extended Data Fig. 9). These effects are partially transcriptional: RNAseq analysis revealed that glycine increases the expression of several enzymes that regulate glycolysis and oxidative phosphorylation (Extended Data Fig. 10, Extended Data Tables 5-7). In cells supplemented with glycine, several enzymes that regulate lipid metabolism and free fatty acid (FFA) oxidation were also upregulated; the latter finding suggested that cyanide may induce a shift towards FFA utilization. Thus, we have tested the effect of etomoxir (a pharmacological agent which inhibits carnitine palmitoyltransferase-1 on the outer mitochondrial membrane and thus suppresses the feeding of FAO-derived acetyl-coenzyme-A into the Krebs cycle) on the bioenergetic profile of HepG2 cells. Etomoxir exerted a more pronounced inhibitory effect on mitochondrial respiration in glycine-treated cells than in control cells and attenuated the stimulatory effect of glycine on mitochondrial respiration (Fig. **4d**, Extended Data Fig. 8e,f).

Cell proliferation is closely tied to cellular ATP generation. We found that exogenously administered cyanide exerted a bell-shaped effect on HepG2 cell proliferation, increasing proliferation at low (nanomolar) concentrations and reducing proliferation at higher (10 micromolar and above) concentrations (Fig. **4e**). In line with these observations, cyanide scavengers decreased, while glycine increased cell proliferation; this latter effect was suppressed by cyanide scavengers (Fig. **4f,g**). Conversely, inhibition of cyanide production, either via lysosomal alkalization or inhibition of peroxidase activity decreased cell proliferation (Fig. **4h,i**). Importantly, both TST overexpression and TST silencing decreased cell proliferation (Fig. **4j**), consistently with the concept that an optimal, physiological range of cyanide level is necessary to support cell proliferation, and significant deviations in either direction (Fig. **2o**) impair bioenergetic function and inhibit cell proliferation.

## Cyanide exerts cytoprotective and organ-protective effects in hypoxia

Physiological concentrations of the known gasotransmitters exert cytoprotective effects**^1–7^**. We, therefore, hypothesized that endogenously generated cyanide may also play such a role. Glycine supplementation protected HepG2 cells from hypoxia and hypoxia/reoxygenation-induced cell death (Fig. **5a**). A low concentration of KCN (10 nM) recapitulated this protection (Fig. **5a**). On the other hand, cyanide scavenging (THC, CoE) or omission of glycine/serine from the culture medium exacerbated hypoxia and hypoxia/reoxygenation induced cell death (Fig. **5a**). Glycine supplementation under normoxia upregulated several genes involved in oxidative stress response (Extended Data Fig. 10, Extended Data Table 8), while in hypoxic conditions glycine attenuated the upregulation of HIF-1α expression (Fig. **5b**). Likewise, significant changes in gene expression were observed both with TST overexpression and TST silencing, including effects on multiple key biochemical pathways relevant for cell metabolism and proliferation and viability (Extended Data Table 9,10).

**Figure 5.**
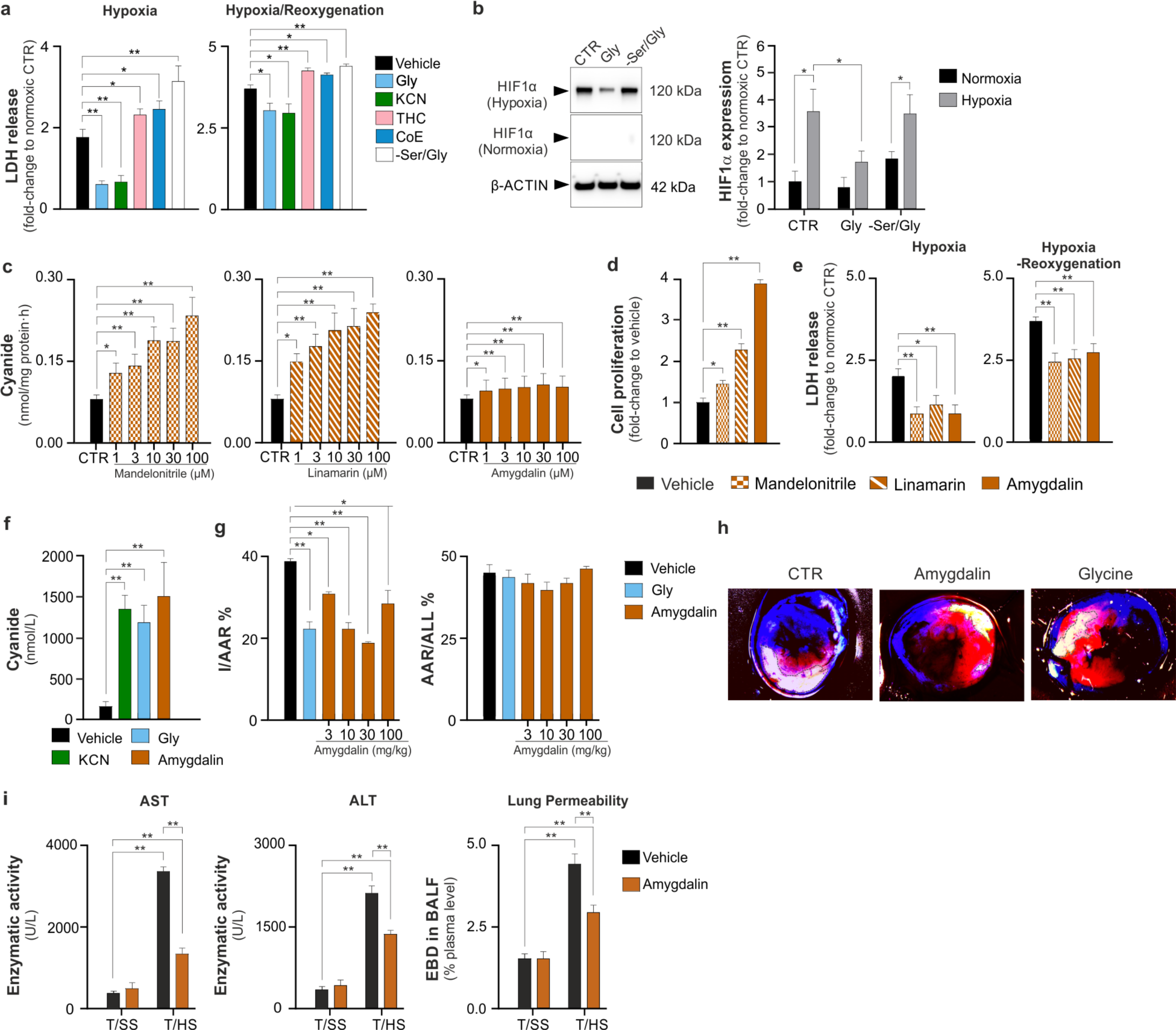
Controlled cyanide supplementation exerts cytoprotective effects. **(a-b)** Effect of treatment with glycine (Gly, 10 mM), authentic cyanide (KCN, 10 nM), cyanide scavengers trihistidyl-cobinamide (THC, 10 µM) or cobalt edetate (CoE, 10 µM) or serine/glycine deprivation (-Ser/Gly) in HepG2 cells subjected to hypoxia (48 h) or hypoxia/reoxygenation (48/24 h) on **(a)** lactate dehydrogenase (LDH) release and **(b)** hypoxia inducible factor 1 subunit alpha (HIF1α) expression. **(c)** Cyanide release from mandelonitrile, linamarin and amygdalin (ECh method). **(d)** Effect of mandelonitrile, linamarin and amygdalin (300 µM) on cell proliferation and **(e)** hypoxia-induced cell injury (LDH release) in HepG2 cells. **(f)** Cyanide concentrations in mouse blood under baseline conditions and after administration of KCN (0.1 mg/kg), glycine (Gly, 100 mg/kg) or amygdalin (10 mg/kg). **(g,h)** Effect of various doses of amygdalin (3-300 mg/kg) or glycine (300 mg/kg) on infarct size in a mouse model of myocardial ischemia/reperfusion. The ischemic area (I) relative to the area at risk (AAR) and AAR relative to the whole (ALL) myocardial area is shown. **(i)** Effect of amygdalin (10 mg/kg) on indices of organ damage (aspartate aminotransferase – AST, alanine transaminase – ALT and lung permeability) in a mouse model of hemorrhagic shock. T/SS: sham-shock; T/HS: hemorrhagic shock. Data are presented as mean±SEM of at least n=5 independent experiments. *p<0.05 or **p<0.01 indicate significant differences.

Similar to low concentrations of exogenous KCN**^30^**, treatment with the cyanogenic compounds mandelonitrile, linamarin and amygdalin (Fig. **5c**) stimulated cell proliferation (Fig. **5d**), recapitulated the cytoprotective effect of glycine and cyanide in hypoxia and hypoxia/reoxygenation (Fig. **5e**), and increased protein cyanylation (Extended Data Fig. 6e,f).

Cyanide concentration in the mouse blood was quantified as 163±36 nM; that administration of amygdalin (10 mg/kg) or glycine (100 mg/kg) to mice increased the blood cyanide concentration 5-fold to ∼1 µM. A comparable increase in the blood cyanide concentration could be achieved by administration of a low, non-toxic dose (0.1 mg/kg) of KCN (Fig. **5f**). Using the same detection method employed in the current study, the cyanide concentration in the blood of healthy non-smoker human volunteers was quantified as 540±10 nM (n=45) **^31^**.

In a mouse model of myocardial ischemia/reperfusion, amygdalin administration and glycine supplementation reduced infarct size (Fig. **5g,h**). In the same model, amygdalin also exerted a protective effect, and exhibited a bell-shaped dose-response, with the highest protective effects obtained at 10 and 30 mg/kg doses. Similarly, in a mouse model of hemorrhagic shock, amygdalin reduced the degree of hepatic and pulmonary injury (Fig. **5i**).

## Severe overproduction of endogenous cyanide induces impairment of cellular bioenergetics in nonketotic hyperglycinemia

Glycine encephalopathy (also known as nonketotic hyperglycinemia or NKH)**^32^** is a devastating disease caused by mutations in the genes *GLDC* or *AMT* (genes that encode essential proteins of the glycine cleavage enzyme system), leading to a pathological buildup of glycine in the cells and blood of NKH patients. We hypothesized that NKH could also result in the cellular accumulation of endogenous intracellular cyanide, potentially contributing to cytotoxic effects. Confocal microscopy revealed that fibroblasts derived from NKH patients – in the standard culture medium containing 400 µM glycine – showed a markedly elevated cyanide signal compared to control fibroblasts (Fig. **6a**). The cyanide signal was strongest in the lysosomes, but was also markedly distributed throughout the cytosol of the NKH cells (Fig. **6b**). As expected, intracellular glycine levels were markedly higher in NKH cells than in normal control fibroblasts from healthy individuals (Fig. **6c**). Quantification of cyanide production by the electrode method revealed that NKH cells generate approximately 30-fold higher rate than control fibroblasts (Fig. **6d**) and treatment with the lysosomal alkalinizer hydroxychloroquine (Hcq) or the cyanide scavenger THC reduced cyanide levels in the NKH cells (Fig. **6d**). Mitochondrial electron transport chain activity, ATP generation (Fig. **6e**), cell viability and proliferation rate (Fig. **6f,g**) were significantly lower in NKH fibroblasts than control cells. Hydroxychloroquine improved the bioenergetics, viability and the proliferation rate of NKH fibroblasts (Fig. **6f-h**). These findings demonstrate that endogenous cyanide production is in NKH is elevated to cytotoxic levels (Fig. **6i**).

**Figure 6.**
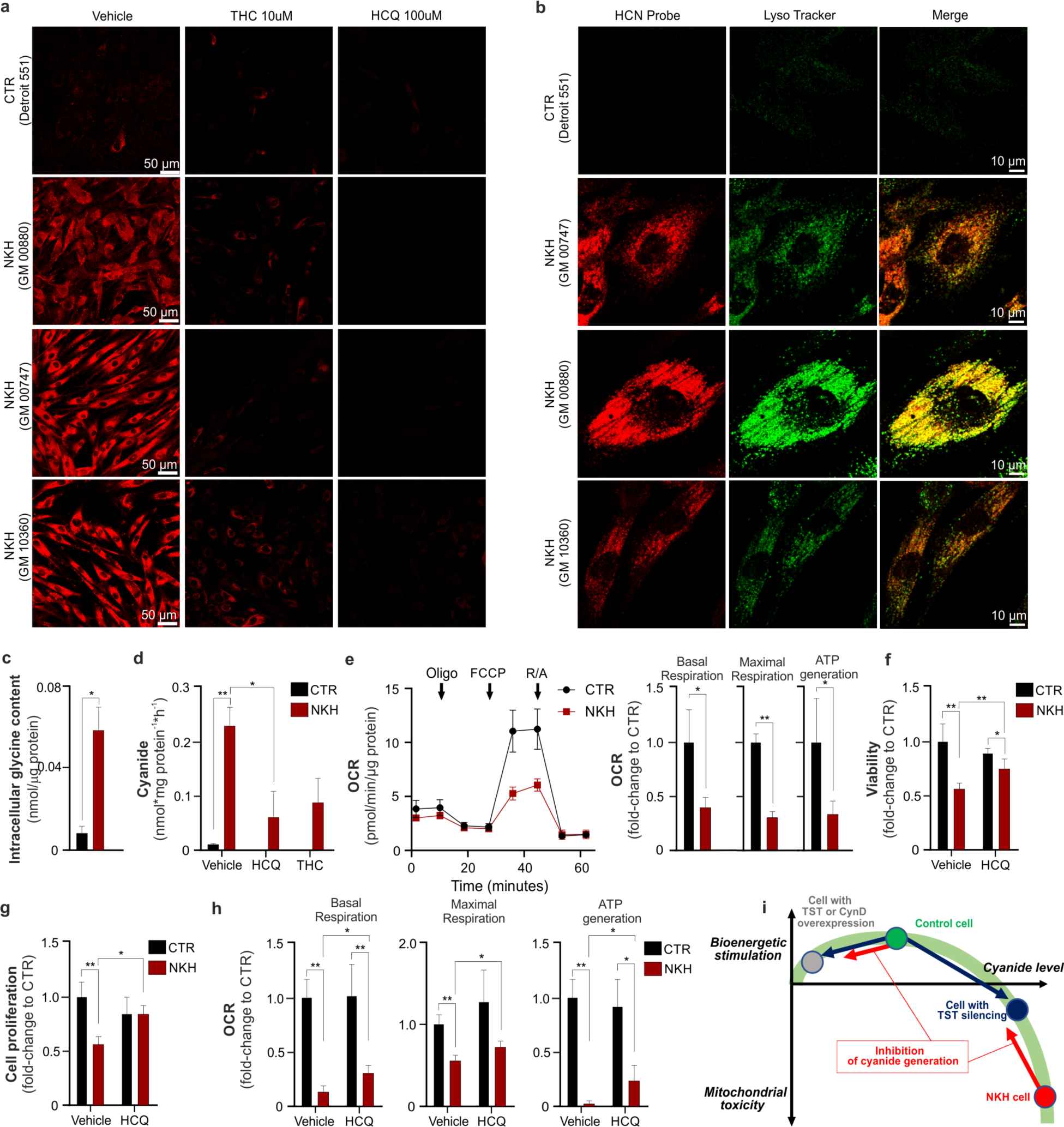
Endogenous cyanide generation is increased and cell function is diminished in fibroblasts derived from nonketotic hyperglycinemia (NKH) patients. **(a)** Confocal microscopy images showing increased endogenous cyanide levels in fibroblasts derived from NKH patients (cell lines GM00880, GM00747, and GM10360) compared to control fibroblasts from healthy individuals under control conditions (Vehicle), in the presence of the cyanide scavenger trihistidyl-cobinamide (THC, 10 µM) or the lysosomal deacidification agent hydroxychloroquine (HCQ, 100 µM). The fluorescence intensity of CSP cyanide selective probe indicates a significant difference between NKH-derived and control fibroblasts. **(b)** Confocal microscopy images showing the partial co-localization of cyanide with lysosomes (LysoTracker) in in fibroblasts derived from NKH patients (cell lines GM00880, GM00747, and GM10360) compared to control fibroblasts from healthy individuals. **(c)** Quantification of intracellular glycine concentration in control (CTR) and NKH fibroblasts. **(d)** Measurement of cyanide production in in control (CTR) and NKH fibroblasts, measured by the electrochemical detection method. Treatment for 24h with the lysosomal alkalinizer hydroxychloroquine (Hcq, 10 µM) or the cyanide scavenger THC (10 µM) reduced cyanide levels in NKH-derived fibroblasts. **(e)** Mitochondrial electron transport chain activity, measured by Extracellular Flux analysis, was significantly reduced in NKH-derived fibroblasts compared to controls, indicating mitochondrial dysfunction. OCR: oxygen consumption rate. Arrows represent the times of the addition of the ATP synthase inhibitor oligomycin, the uncoupling agent carbonyl cyanide-4-(trifluoromethoxy)phenylhydrazone (FCCP) and the combined addition of the mitochondrial Complex I inhibitor rotenone and the mitochondrial Complex III inhibitor antimycin (R/A) in the Extracellular Flux Analysis protocol. **(f, g)** Cellular viability (LDH release) and proliferation rate (BrdU incorporation) in fibroblasts derived from NKH patients were significantly lower than in control fibroblasts; treatment of the cells with hydroxychloroquine (HCQ, 30 µM, 72h) improved these parameters. **(h)** Cellular bioenergetic parameters in NKH cells after 72h treatment with hydroxychloroquine (HCQ, 30 µM). **(i)** The bell-shaped concentration-response curve of cyanide in mammalian cells: at physiological concentrations, cyanide supports mitochondrial function, stimulates metabolism and contributes to the maintenance of proliferation. Rhodanese (thiosulfate sulfurtransferase, TST) or cyanide dihydratase (CynD) overexpression results in an increased decomposition of endogenously generated cyanide, and attenuates these stimulatory effects; similar mechanism is responsible for the bioenergetic effect of cyanide scavengers or inhibitors of cyanide generation in healthy control cells (red arrow). TST silencing attenuates the decomposition of endogenously generated cyanide; cyanide accumulates and reaches levels at which it impairs mitochondrial function and suppresses bioenergetics and proliferation. When cyanide is generated at very high rates (such as in NKH cells where the glycine cleavage system is impaired), cyanide reaches concentrations where it markedly impairs metabolism and proliferation and exerts cytotoxic effects. Inhibition of cyanide generation in NKH cells attenuates these toxic effects (red arrow). Data are presented as mean±SEM of at least n=5 independent experiments. *p<0.05 or **p<0.01 indicate significant differences.

## Discussion

Cyanide is generally considered a toxic molecule to mammalian cells due to its ability to inhibit mitochondrial respiration at Complex IV**^8^**. However, recent studies indicate that KCN administration to mammalian cells, at nanomolar to low micromolar concentrations stimulates mitochondrial respiration, promotes cell proliferation, and protects the cells from oxidative damage**^30,33^**. While these data predicted that cyanide may have regulatory roles in mammalian cells, the question of whether cyanide is an *endogenous* mammalian gaseous mediator remained to be addressed. Cyanide is known to play such roles in certain bacteria and plants**^34,35^**. The current report demonstrates that mammalian liver and spleen produce cyanide under basal conditions, and that cyanide generation can be increased with glycine supplementation. Biochemical assays demonstrated that cyanide production *in vitro* occurs with at a pH optimum of 4.5, a pH typically found only in lysosomes. Indeed, lysosomes were shown to produce cyanide *in situ* as well as *ex vivo* and cyanide generation was stimulated by adding glycine. Conversely, disrupting the lysosomal pH or inhibiting the lysosomal glycine transporter reduced cyanide production. Similar to what was recently shown for low concentrations of exogenously supplied KCN**^30^**, we conclude that endogenously produced cyanide in mammalian cells and tissues exerts physiological roles such as (i) support of mitochondrial respiration, (ii) stimulation of cell proliferation and (iii) cytoprotection. These roles appear to be applicable for various parenchymal cells; while the current project focused on hepatocytes, detectable cyanide generation was also found in cultured hepatocytes, epithelial cells, endothelial cells, fibroblasts as well. Moreover, cells of monocytic and neutrophilic lineage (U937, HL60) and primary human PBMCs and neutrophils also produced high amounts of cyanide, both basally and after incubation with glycine. The role of cyanide generation in the function of immune cells is intriguing and remains a subject of further investigation.

The mechanism of cyanide’s action in mammalian cells is likely multifaceted. In the current study we focused on the cyanylation of protein cysteine residues. Cyanylated proteins have previously been detected in plants and human plasma and sub-micromolar concentrations of cyanide have been found in human blood**^28,29,31,36^**. The current report shows that hundreds of proteins are physiologically cyanylated in mammalian cells and that the portion of the proteins affected by this modification could be quite high. Based on the experiments with GAPDH and GPDH, the functional response to cyanylation can be either stimulatory or inhibitory, depending on the particular protein. This is similar to posttranslational SH group modifications elicited by other gasotransmitters – nitrosylation by NO, persulfidation by H_2_S – which can be stimulatory or inhibitory, dependent on the particular protein**^6,7^**. Among the proteins whose cyanylation was found to increase upon glycine supplementation (Extended Data Table 1,2), acyl-CoA synthetase short-chain family member 3 (mitochondrial), 3-ketoacyl-CoA thiolase A (peroxisomal), acetyl-CoA synthetase (cytoplasmic), long-chain specific acyl-CoA dehydrogenase (mitochondrial), and glycerol-3-phosphate dehydrogenase (mitochondrial) are of particular interest. These proteins are directly involved in beta-oxidation and fatty acid biosynthesis. We propose that endogenous cyanide – through a combination of transcriptional and post-translational mechanisms – acts as a global regulator of cell function. One aspect of this re-programming is a shift towards oxidation of FFAs, which likely occurs through a combination of translational and posttranslational effects of cyanide at multiple enzymes that regulate the catabolism of FFAs. This type of shift is known to occur physiologically, for instance, in response to fasting or prolonged exercise**^37^**. Also noteworthy are the observed cyanylation effects (Extended Data Table 1,2) on enzymes in the transsulfuration pathway and glutathione biosynthesis, such as glutamate-cysteine ligase (catalytic subunit), betaine-homocysteine S-methyltransferase 1, and cystathionine gamma-lyase (CSE). The latter enzyme is of the key enzymes involved in transsulfuration and H_2_S biosynthesis**^6^**. Cyanylation of CSE could provide a potential link for cross-talk of cyanide signaling with H_2_S signaling, which remains to be addressed in future studies.

Glycine is a common, non-essential amino acid that is generated endogenously in mammalian cells through *de novo* synthesis, and is supplied by dietary sources. It enters cells from the extracellular space through various transporters, and is used for the *novo* synthesis of proteins and nucleotides**^38^**. Its intracellular concentration is in the 3-10 mM range, while its lysosomal concentration is ∼0.5 mM**^38–41^**. Glycine supplementation is cytoprotective in various models of hypoxia and ischemia and reperfusion *in vitro* and *in vivo***^41^**. Based on the current data, we propose that part of the beneficial effect of glycine may be due to its ability to stimulate the synthesis of low – cytoprotective – concentrations of cyanide; direct measurements of blood cyanide concentration show that 100 mg/kg glycine produces a peak blood cyanide concentration of approximately 1,200 nM. Thus, at the therapeutically effective doses of glycine, blood cyanide concentration likely remains in the single-digit micromolar concentration range. We *do not propose* that *slight changes* in extracellular glycine can affect cellular cyanide generation, because of the high concentrations of intracellular glycine in mammalian cells. The fact that modest changes in the extracellular concentration of an amino acid precursor do not affect the generation of various gasotransmitters in healthy normal cells is well known in the biology of NO and H_2_S. Although L-arginine is the precursor of NO, addition of extracellular L-arginine in most cases does not stimulate NO production. Likewise, H_2_S is produced from cysteine and homocysteine by mammalian cells, but addition of these molecules to cells does not drive additional H_2_S generation in most normal cells and tissues**^1–7^**.

Cytoprotective concentrations of cyanide may also be generated through administration of low concentrations of cyanogenic molecules – as exemplified in the current report by amygdalin, mandelonitrile and linamarin. These molecules are primarily known for their potential toxicity due to cyanide release; they have been tried as cytotoxic agents as a treatment for cancer, an approach that is severely hampered by toxicity to the host.**^42^** In sharp contrast to this prior concept, the current data suggest that administration of low amounts of cyanogenic compounds could be an experimental therapeutic strategy. When mandelonitrile, amygdalin or amygdalin are added to HepG2 cells at a concentration range of 1-100 µM, the cyanide generation rates are in the range of 0.1-0.2 nmoles/mg/protein/h, which is comparable to the endogenous generation rates of cyanide in these cells. Based on direct blood measurements (Fig. **5f**), we estimate that cyanide blood concentrations in the optimal therapeutic dose range of amygdalin (10-30 mg/kg) are in the 200-1,000 nM range *in vivo*.

The bell-shaped dose-response effect, a fundamental characteristic of gasotransmitter biology, emphasizes the balance between cytoprotective and cytotoxic effects of gasotransmitters based on their concentration. This balance depends on the cell’s capacity to produce and detoxify these compounds, and once this capacity is exceeded, the result is often detrimental to cellular health**^1–7^**. Indeed, the current and prior findings**^30,33,34^** demonstrate the bell-shaped concentration-response of exogenously administered cyanide on cellular bioenergetics and proliferation, and a similar bell-shaped concentration-response curve applies for endogenously generated cyanide as well (Fig. **6i**). A bell-shaped concentration-response was also evident for amygdalin in our myocardial infarction model (Fig. **5g**); similar bell-shaped concentration-responses have been previously observed for NO or H_2_S donation in this model as well**^43,44^**. In fibroblasts from NKH patients the pathological accumulation of glycine leads to markedly increased (approximately 30-fold) cyanide production rates, crossing the cytotoxic threshold. The improved bioenergetic function and viability of NKH-derived fibroblasts following treatment with the lysosomal alkalinizer hydroxychloroquine (Hcq) indicate that lysosomal generation of cyanide plays a role in the observed cellular dysfunction. The marked reduction in mitochondrial electron transport chain activity and cellular proliferation rates in fibroblasts from NKH patients further support this view (Fig. **6i**). These findings could have implications for understanding the pathophysiology of glycine encephalopathy and potentially offer new therapeutic approaches for managing this condition.

Although most of the scientific literature related to the role of cyanide in mammals focuses on its toxicological properties, one should emphasize that cyanide is generated endogenously in several bacterial and plant species, serving various regulatory effects, such as quorum sensing, biocontrol, germination development, and immunity**^35,45,46^**. Indeed, cyanide – together with H_2_S – were already present on the planet several billions of years ago, before the appearance of atmospheric oxygen, and bacteria and plants have been linked to early biochemical processes that led to evolution of higher organisms**^47–51^**. In this context, it makes sense that nature developed the enzymatic machinery to produce these gases despite their obvious toxicity. Likewise, their important reactions, particularly with protein cysteine residues**^52–54^**, play evolutionally conserved roles and functions across the animal and plant kingdom. In this context, the current findings place cyanide – together with NO, CO and H_2_S – in the group of mammalian regulatory gasotransmitter species (Extended Data Table 11). These gases play various regulatory roles, which are sometimes distinct, sometimes overlapping, and sometimes cooperative**^1,6,52–56^**. Further studies remain to be conducted to further characterize the role of endogenous cyanide in mammalian cells and tissues. These studies will focus on its potential interactions with various other gaseous mediators, as well as the role of the glycine/cyanide system in healthy cells and in various diseases.

## Methods

### Animals

C57BL/6J mice were purchased from Janvier Laboratories (Le Genest-Saint-Isle, France). Myeloperoxidase knockout mice (Mpo^-/-^ strain: #004265) and peroxidasin heterozygous mice (Pxdn^+/-^; strain: #042166), both on C57BL/6J background, were purchased from Jackson Laboratories. Despite repeated breeding efforts, we were unable to generate Pxdn^-/-^ mice, and therefore tissues from Pxdn^+/-^ mice were used. Animals were housed in a light-controlled room with a 12 h light-dark cycle and were allowed ad libitum access to food and water. All studies were performed on 12-18 weeks-old male mice.

### Myocardial ischemia/reperfusion injury model

Myocardial infarction was induced by ligation of the left coronary artery as previously described**^57^**. Male 8–10-week-old C57BL/6J mice were randomly divided into 6 groups. Control group (n=10) received saline vehicle only. Amygdalin (3, or 10, or 30 or 100 mg/kg; Cayman, 26668) was administered intraperitoneally at a dose of 10 mg/kg, 1 hour before ischemia (n=7). Glycine (300 mg/kg, Fisher Scientific, BP381) was administered intraperitoneally at a dose of 300 mg/kg, 1 h before ischemia (n=9). Animals were anesthetized by intraperitoneal injection with a combination of ketamine and xylazine (0.01 ml/g, final concentrations of ketamine, xylazine 10 mg/ml and 2 mg/ml, respectively). Anesthetic depth was evaluated by the loss of pedal reflex to toe-pinch stimulus and breathing rate. Additional anesthesia (1/4 of the dose) was applied during the first hour of reperfusion to maintain the anesthesia depth. A tracheotomy was performed for artificial respiration at 150 strokes/min with a tidal volume of 200 μl. A thoracotomy was then performed, and the pericardium was carefully retracted to visualize the left anterior descending (LAD) coronary, which was ligated using a 6-0 silk suture (Ethicon, W888) placed 3 mm below the tip of the left atrium with the help of 5 mm piece of a 1 mm diameter catheter tube. The heart was allowed to stabilize for 15 minutes prior to ligation to induce ischemia. After the ischemic period, the ligature was released and allowed reperfusion of the myocardium. Throughout the surgical procedure, body temperature was maintained at 37 + 0.5 °C with a heating pad. After reperfusion, hearts were rapidly excised from mice and directly cannulated through aorta and washed with Krebs buffer (NaCl 118.5 mM, NaHCO_3_ 25 mM, KCl 4.7 mM, MgSO_4_ 1.2 mM, KH_2_PO_4_ 1.2 mM, glucose 11 mM and CaCl_2_ 1.5 mM, pH=7.4) for blood removal. Hearts were perfused with 500 μl 2% Evans Blue, diluted in Krebs buffer. Afterwards they were kept at -80 °C for 1 h and then sliced in 2 mm sections parallel to the atrioventricular groove. The slices were incubated in 2 ml of 1% TTC phosphate buffer (PBS pH =7.4) at 37 °C for 10 min. Slices were then compressed between glass plates 1 mm apart and photographed with a Leica DFC310 FX Digital Color Camera through a Nikon SMZ800 stereoscope and measured with the NIH ImageJ. Measurements were performed in a blinded fashion. The myocardial area at risk as well as the infarcted and the total area were automatically transformed into volumes. Infarct and risk area volumes were expressed in cm^3^ and the percentage of infarct-to-risk area ratio (%I/AAR) and of area at risk to whole myocardial area (% AAR/All) were calculated as described**^57^**.

### Hemorrhagic shock model

C57BL/6J mice were randomly assigned into the following groups: trauma/sham shock (T/SS) receiving vehicle, T/SS receiving amygdalin, trauma/hemorrhagic shock (T/HS) receiving vehicle, and T/HS receiving amygdalin. In these pharmacological experiments the mice were pretreated (30 minutes before T/SS or T/HS) intraperitoneally either with vehicle or amygdalin (10 mg/kg). Mice were anesthetized with 1% isoflurane and rectal temperature was maintained between 36.5 and 37.5 °C using a feedback-controlled homeothermic blanket heating system (Sumno-suite, Kent Scientific). A fixed-pressure model was used to induce hemorrhagic shock. Briefly, after anesthesia mice received a midline laparotomy of 2 cm and then the incision was closed with 4-0 silk suture (Covetrus, 034902). The right and left femoral arteries were isolated and catheters were placed for monitoring blood pressure and blood withdrawal, respectively. For blood withdrawal, a sterile 1-ml syringe with a 30G needle, which was attached to PE-10 tubing and filled with 0.2 ml of 1% heparinized saline was used. Each mouse received 1U heparin. Blood pressure was monitored using a continuous blood-pressure monitoring system (Powerlab 8/30, AD Instruments, Colorado Springs, CO, USA). After five minutes of baseline recording mice were treated with a drug or vehicle, which was followed by inducing shock for a 2.5 h period. Blood pressure was maintained between 28-32 mmHg by withdrawing or reinfusing the shed blood. At the end of the shock period, mice were resuscitated with Ringer’s Lactate at three times the amount of shed blood for 15 minutes. Three hours after resuscitation the mice were euthanized and BALF, blood and tissues were collected as described below. Blood pressure was analyzed using LabChart software (AD Instruments, Colorado Springs, CO, USA). T/SS animals were exposed to the same procedures except for blood withdrawal**^58^**.

The liver enzymes aspartate aminotransferase (AST) and alanine aminotransferase (ALT) levels were determined from plasma samples. Plasma samples were diluted (1:10) with AST (Thermo Fisher, TR70121) and ALT (Thermo Fisher, TR71121) reagents and the data were acquired using a spectrophotometer at 340 and 405 nm. Three hours after the T/HS period, mice were re-anesthetized with isoflurane. Evans blue dye (EBD, Sigma-Aldrich, E2129) was administered through the tail vein and five minutes later about 1 ml of blood was withdrawn from tail artery. Twenty minutes later the mice were euthanized and the trachea was isolated for bronchoalveolar lavage fluid (BALF) sample collection. After a small incision, a syringe with a 23G needle filled with 1 ml of sterile saline was placed in the trachea. Lungs were injected and aspirated two times and the BALF was collected. To measure the EBD in BALF, the BALF sample was centrifuged at 4 °C at 1,500 x g for 20 minutes. The supernatant was collected and assayed at 620 nm spectrophotometrically. The concentration of Evans blue dye in the BALF was then expressed as a percentage of its plasma concentration**^58^**.

### Cell culture

The HepG2 hepatocellular carcinoma cell line (ATCC^®^ HB-8065^TM^), CynD-over-expressing HepG2 cells, TST-over-expressing HepG2 cells and TST-knockdown (shTST) HepG2 cells were grown in Dulbecco’s Modified Eagle Medium (DMEM) culture medium containing 1.0 g/l D-glucose (Gibco, 21885), supplemented with 10% (v/v) heat-inactivated fetal bovine serum FBS (Hyclone), 100 units/ml of penicillin and 100 µg/ml of streptomycin. The Hep3B hepatocellular carcinoma cell line (ATCC^®^ HB-8064^TM^) and HL-60 cells (ATCC^®^ CCL-240) were grown in DMEM culture medium containing 1.0 g/l D-glucose (Gibco, 21885), supplemented with 10% (v/v) heat-inactivated FBS (Hyclone), 100 units/ml of penicillin and 100 µg/ml of streptomycin. Human umbilical vein endothelial cells (HUVECs) were grown in Endothelial Cell Growth Medium (211-500, Cell Applications Inc., San Diego, CA, USA) The HEK293A human embryonic kidney cell line (kind gift from Chick Wallace), MPO-over-expressing HEK293T cells, PXDN-over-expressing HEK293T cells and catalase-over-expressing HEK293T cells were cultured in DMEM containing 4.5 g/l glucose (Gibco, 11965). The culture medium was supplemented with 10% (v/v) heat-inactivated FBS (Hyclone), 2 mM Glutamax, non-essential amino acids, 100 units/ml penicillin and 100 μg/ml streptomycin. The A549 lung carcinoma epithelial cell line (ATCC^®^ CCL-185^TM^) was cultured in DMEM 4.5g/l D-glucose (PAN Biotech, P04-03500). The culture medium was supplemented with 10% (v/v) heat-inactivated FBS (Hyclone), 100 units/ml penicillin and 100 μg/ml streptomycin. The HCT116 colorectal carcinoma cell line (ATCC^®^ CCL-247^TM^) and the HT29 colorectal adenocarcinoma cell line (ATCC^®^ HTB-38^TM^) were cultured in McCoy’s 5A Medium (Gibco, 16600). The culture medium was supplemented with 10% (v/v) heat-inactivated FBS (Hyclone), 100 units/ml penicillin and 100 μg/ml streptomycin. LoVo human colorectal adenocarcinoma cell line (ATCC^®^ CCL-229^TM^) was cultured in Advanced Dulbecco’s Modified Eagle Medium/nutrient mixture F-12 (DMEM/F-12, 1:1, 1X) (Gibco, 12634). The culture medium was supplemented with 10% (v/v) heat-inactivated FBS (Hyclone), 100 units/ml penicillin and 100 μg/ml streptomycin. The U937 pro-monocytic, human myeloid leukemia cell line (ATCC^®^ CRL-1593.2^TM^) was cultured in RPMI-1640 Medium (ATCC 30-2001). The culture medium was supplemented with 10% (v/v) heat-inactivated FBS (Hyclone), 2 mM Glutamax, 100 units/ml penicillin and 100 μg/ml streptomycin. Human dermal fibroblasts from a healthy subject (Detroit551, ATCC^®^ CCL-110^TM^) were cultured in Advanced Dulbecco’s Modified Eagle Medium/nutrient mixture F-12 (DMEM/F-12, 1:1, 1X) (Gibco, 11320). The culture medium was supplemented with 0.1% lactalbumin hydrolysate, 10% (v/v) heat-inactivated FBS (Hyclone), 100 units/ml penicillin and 100 μg/ml streptomycin. The U138-MG human glioblastoma cell line (ATCC^®^ HTB-16™) was cultured in DMEM (ATCC 30-2002). The culture medium was supplemented with 10% (v/v) heat-inactivated FBS (Hyclone), 100 units/ml penicillin and 100 μg/ml streptomycin. Serine/glycine free medium was obtained from USBiological (D9802-01) supplemented with 4.5 g/l D-glucose, 2 mM glutamine, 1 mM sodium pyruvate, 10% (v/v) heat-inactivated FBS (Hyclone), 100 units/ml penicillin and 100 μg/ml streptomycin. Human primary hepatocytes (from a 48-year-old Caucasian male) were purchased from AnaBios Corporation (San Diego, CA, USA). Cryopreserved cells were thawed in HEP-005 Anabios thawing medium, plated in HEP-003 Anabios plating medium and maintained in HEP-004 Anabios maintenance medium. Human dermal fibroblasts from NKH patients GM00880 (from a 21-year-old Caucasian male), GM00747 (from a 1.5 years-old Caucasian female), GM10360 (from a 2-months-old Caucasian male), were obtained from Coriell Institute for Medical Research (Camden, NJ, USA). Cells were cultured in DMEM (Hyclone, SH30243.01), supplemented with 15% (v/v) heat-inactivated FBS (Hyclone), 1% non essential aminoacids (Hyclone) and 100 units/ml penicillin and 100 μg/ml streptomycin.

All cells were grown in a humidified incubator at 37 °C and 5% CO_2_ atmosphere. For experiments and sub-culturing, cells were rinsed with PBS and detached from T75 flasks by incubating with 0.25% (w/v) Trypsin- 0.53 mM EDTA for 2-5 min at 37 °C followed by resuspension in culture medium.

### Generation of stably transfected cell lines

Lentivirus Gene Expression Vectors: The lentivirus gene expression vector pLV[Exp]-Bsd-CMV was obtained from VectorBuilder (Chicago, IL, USA). The human genes: MPO (coding myeloperoxidase protein); PXDN (coding peroxidasin protein); CAT (coding catalase protein); CynD (coding cyanide dehydratase protein) and TST (coding thiosulfate sulfurtransferase protein) were codon optimized and contained a Myc tag. The gene synthesis and the cloning of the genes to pLV vector were performed by GeneScript (NJ, USA).

*HepG2 ShTST generation:* HEK293 T cells (ATCC, American Type Culture Collection, Manassas, VA, USA) were seeded at 70–80 % confluence in a 6-well plate and transiently transfected 4-6 h later with the TST shRNA plasmid (sc-36418-SH : Santa Cruz Biotechnology Inc., Santa Cruz, CA 95060), a packaging plasmid pLP1, pLP2, and an envelope plasmid pLP/ VSVG in a ratio of 4.2:2:2.8 using JetOptimus (Polyplus, Strasbourg, France) according to the manufacturer’s instructions. The transfection mixture and medium were replaced with fresh culture medium after o/n incubation. Lentiviral supernatants were collected after 24 h and filtered via a 0.45 μm filtration unit and subsequently aliquoted and stored at -80 °C until use. Human HepG2 cells were seeded in 12-well plates to reach approximately 50 % confluence on the day of transduction. The cells were transduced with lentiviral supernatant in the presence of 6 μg/mL protamine sulfate. Fresh complete DMEM medium was added 24 h after transduction. Seventy-two hours following transduction, puromycin (2 μg/ml) was added to the culture to select transduced cells for 3 days.

*Lentiviral production and cell transduction*: Viral particles were produced in HEK293T cells using a third generation lentiviral system. Lentiviral supernatants were collected after 24 h and filtered with 0.45 μm filtration unit and subsequently aliquoted and stored at -80 °C until use. Human HepG2 cells and human HEK293A were transduced with lentiviral supernatant in the presence of 6 μg/ml protamine sulfate. Seventy-two hours following transduction, 5 μg/ml or 45 μg/ml Blasticidin S (InvivoGen, Toulouse France) was added to the culture to select transduced cells.

### HCN detection

KCN and its solutions are poisonous to humans; they were handled with care in a well-ventilated hood. Additionally, HCN is released from solutions with pH values near or below the pKa of HCN (pKa = 9.2). Thus, all aqueous standards containing cyanide were prepared from KCN in NaOH (10 mM or 0.5 M, depending on the detection method) to ensure that cyanide remains in its non-volatile form (CN^-^).

*Sample preparation, mouse blood:* Mice were divided into a control group 0.9 % NaCl (n=10), a potassium cyanide (KCN) group (0.1 mg/kg, n=7), a glycine group (100 mg/kg, n=7) and an amygdalin group (10 mg/kg, n=7). KCN, glycine or amygdalin were diluted in water for injection and administered intraperitoneally. The dose of KCN used in this study was over 50-times lower than the LD_50_ and 10 times lower than the LOAEL (lowest observable adverse effect) dose in mice. After 10 min (KCN) or 3 h (glycine or amygdalin), mice were euthanized using the intraperitoneal injection of ketamine (100 mg/kg) and xylazine (10 mg/ml) and heart blood was collected after 5 min in heparinized and airtight tubes closed with screw caps. 200 µl of blood was immediately frozen at -80 °C, in a tightly sealed tube and stored for further analysis with the Cyanalyzer LC-MS/MS method (see below).

*Sample preparation, mouse tissue*: Mice were sacrificed using CO_2_ and exsanguination. The animals were perfused using 20 ml chilled PBS through the ascendent aorta for 2-4 min to remove blood from the tissues. Tissues were surgically removed (liver, without gallbladder) and placed on ice. 20-30 mg of tissues (liver, kidney, lung, brain, spleen) were placed in a two ml Eppendorf-tube and homogenized in 2 ml PBS using three 2.4 mm metal beads (Omni International, 19-640-3) using a bead mill 4 mini homogenizer (Fisherbrand^TM^) for 240 s, with a speed of 4 m/s. Samples were treated with 10 mM glycine, 10 µM trihistidyl-cobinamide (THC) or 10 µM dicobalt-edetate (Kelocyanor, CoE, US Biological, 446719) and incubated at 37 °C for 24 h. For proteins heat-inactivation, tissue homogenates were incubated at 95 °C for 1 hour. For the physical inactivation of proteins (Freeze-Thawing, F&T), homogenates were subjected to 3 homogenization cycles made of sonication (30 sec pulse followed by 30 sec pause, repeated 5 times using an Ultrasonic Bath Sonicator), freeze (at -20°C for 30 min) and thaw (at 37°C for 1 min). In a separate group of liver homogenates, proteins were denatured by incubation of the tissue homogenates with 2% SDS for 24 h at 37°C. Cyanide production rates were measured with electrochemical, LC-MS/MS and MCC methods (see below).

*Sample preparation: cultured cells:* Cells were seeded in a sterile transparent 6-well plate at 500,000 cells/well (HepG2, Hep3B, HT29, LoVo), 300,000 cells/well (HEK293A, Detroit 551, GM00880, GM00747 and GM10360), 250,000 cells/well (A549), 200,000 cells/well (HCT116), 50,000 cells/well (U138-MG) and 2,000,000 cells/well (U937) and incubated at 37 °C and 5% CO_2_. The day after, medium was replaced with fresh medium, fresh medium containing glycine (10 mM), serine/glycine free medium (-Ser/Gly) or with -Ser/Gly medium supplemented with increasing concentrations of glycine (1, 5 or 10 mM) and cells were incubated at 37 °C and 5% CO_2_ for 24 h (only in the case of cells in suspension, such as U937, cells were seeded directly in - Ser/Gly medium ± glycine). Human primary hepatocytes were seeded at 1,000,000 cells/well in 6-well plate coated with collagen in plating medium. After 6 h, medium was replaced with maintenance medium ± 10 mM glycine and further incubated for 24 h. HCN scavengers were employed to reduce HCN concentrations in cells. For human primary hepatocytes, the day after seeding, cells were treated for 3 h with 10 µM THC. For HepG2 cells, the day after seeding, cells were treated with increasing concentrations of HCN scavengers THC or CoE (1 – 30 µM) and incubated at 37 °C and 5% CO_2_ for 3 h. Incubation (24h) with the glycine-transporter-1 inhibitor iclepertin (LLC HY-138935), at 10-100 µM (Medchemexpress, Monmouth Junction, NJ, USA) was used to inhibit the uptake of glycine into the cells. Increasing concentrations (0 – 100 µM) of the peroxidase inhibitor phloroglucinol (Phl, Sigma-Aldrich, 79330) or a selective MPO inhibitor (AZD-5904, Sigma-Aldrich, SML3274) (24 h) were used to inhibit peroxidase activity. Hydroxychloroquine (Hcq, Sigma-Aldrich, H0915) 1-30 µM (24 h) or bafilomycin (Baf, Alfa Aesar, J61835) 0.01-1 µM (3 h) were used to increase the intralysosomal pH. Glycine-Glycine (Gly-Gly, Sigma-Aldrich, G1002-25G) 150 mM (24 h) was used to inhibit glycine uptake into the cells. The serine hydroxymethyltransferase inhibitor SML2699 (iSHMT, Sigma-Aldrich) (100 µM, 24 h) or lometrexol hydrate (1-10 µM, 24h) (Sigma-Aldrich, SML0040) were used to inhibit serine/glycine intracellular conversion. Various HCN releasers were used to increase HCN levels in HepG2 cells. One day after seeding, cells were treated with increasing concentrations (0 – 100 µM) of the HCN donors amygdalin (Sigma-Aldrich, A6005), linamarin (Toronto, TRCL466000) or mandelonitrile (Sigma-Aldrich, 116025) and incubated at 37 °C and 5% CO_2_ for 24 h. In all cases, after treatment, an aliquot of the supernatant was mixed (1:1, v/v) with 1 M NaOH for HCN measurements using the electrochemical method (see below).

*Sample preparation: primary cells:* Whole blood leukoreduction filters containing total blood leucocytes from the Transfusion CRF Fribourg (Fribourg, Switzerland). PBMCs were isolated from whole blood leucocytes through density gradient centrifugation using Cytiva Ficoll-Paque™ PLUS Medium (Cytiva, Marlborough, Massachusetts, USA). Human Peripheral Blood Neutrophils were purchased from StemCell technologies (Vancouver, British Columbia, Canada). PBMCs or neutrophils were maintained in Gibco™ RPMI 1640 Medium, GlutaMAX™ Supplement (Gibco, Thermo Fisher Scientific, Waltham, MA, USA) supplemented with 10 % FBS (Gibco, Thermo Fisher Scientific), 100 units/mL of penicillin and 100 μg/mL of streptomycin (Gibco, Thermo Fisher Scientific), 2 mM L-Glutamine (Gibco, Thermo Fisher Scientific), 1 mM Sodium Pyruvate (Gibco, Thermo Fisher Scientific), 0.055 mM 2-Mercaptoethanol (55 mM) (Gibco) and 10 mM HEPES (Cytiva). For cyanide production assays, cells were incubated (in closed cryotubes) with PBS or glycine (10 mM) for 3h at 37°C. Then 75ul of the supernatant was collected in a 0.2 ml tube containing 75 µl of NaOH (1 M) and, after incubation at room temperature for 30 minutes, cyanide levels were measured using the cyanide electrode method. The rest of the cells were centrifuged and pellet was obtained. Cell lysates were obtained using 100 µl lysis buffer and BCA measurements were performed to obtain mg of protein to normalize the cyanide production values.

*Cell lysates.* HepG2 over-expressing CynD or TST or knocked down for TST (shTST) were detached with 0.25% (w/v) trypsin- 0.53 mM EDTA for 2-5 min at 37 °C, resuspended with culture medium and collected by centrifugation at 1,000 x g for 5 min. Cell pellets were washed twice with ice-cold PBS and lysed with Cell-Lytic^TM^ M buffer supplemented with 1% phosphatase/protease inhibitor cocktail Halt, 1861281). Lysates were incubated on ice for 30 min and sonicated with 3 cycles of 15 s pulse followed by 15 s pause on ice, using an ultrasonic bath sonicator. Lysates were centrifuged at 17,000 x g at 4 °C for 20 min. The supernatant was collected and total protein were quantified with BCA method (Thermo Scientific™, 23225). For the HCN degradation assay, 500 ng of total protein was incubated in 50 mM Tris-HCl, pH 7.4, in presence of 100 µM KCN in a final volume of 25 µl ± 1 mM sodium thiosulfate (Na_2_S_2_O_3_). HCN degradation activity was stopped at different time-points (0-1 h) by adding 25 µl 1 M NaOH and cyanide detection was performed with electrochemical method (see below).

*Lysosomes.* Lysosome enrichments were performed as described**^59^** from mouse liver or from HepG2 cells. For each experimental day, two mouse livers of wild-type C57BL/6J mice were pooled. Lysosomes were isolated using the lysosome isolation kit (Sigma-Aldrich, MAK405). Tissues were minced and homogenized with a 3 ml glass-teflon Potter-Elvehjem homogenizer (25 strokes) and incubated on ice for 45 min in lysosome isolation buffer. Samples were centrifuged at 500 x g at 4 °C for 10 min and the supernatant was further centrifuged at 20,000 x g at 4°C for 20 min. The resulting supernatant was the cytosolic fraction (referred as Cyto), while the pellet was loaded in a discontinuous density gradient and centrifuged at 147,000 x g at 4°C for 2 h. The layer corresponding to the lysosomal fraction was marked as Lyso, while the other fractions were pooled together and marked as extra-lysosomal fraction (Extra-Lyso). Both lysosomal and extra-lysosomal fractions were resuspended in 200 µl of Suspension Buffer (SB, 10 mM HEPES, pH 7.4, 150 mM NaCl supplemented with phosphatase/protease inhibitor cocktail Halt, 1861281).

For each sample of HepG2 cells lysosomes, cells from 5 × T175 confluent flasks were pooled together. When treatments were performed, cells were treated before collection: DMEM culture medium (CTR); 1 µM Baf, 1 h; 100 µM Hcq, 3 h; 150 mM Gly-Gly, 24 h) and incubated at 37 °C and 5% CO_2_. After treatment, cells were washed with PBS, detached with 0.25% (w/v) Trypsin- 0.53 mM EDTA and resuspended in 10 ml DMEM. Cells were centrifuged at 1,000 x g at 4°C for 5 min. The pellet was washed twice with 10 ml of ice-cold PBS, resuspended in 900 µl of hypotonic buffer 0.1x (HypoB, 3.5 mM Tris-HCl, pH 7.8, 2.5 mM NaCl, 0.5 mM MgCl_2_ supplemented with phosphatase/protease inhibitor cocktail Halt, 1861281) and incubated on ice for 1 h. Cells were homogenized using a 3 ml glass-teflon Potter-Elvehjem homogenizer (100 strokes) and 100 µl of hypertonic buffer 10x (HyperB, 350 mM Tris-HCl, pH 7.8, 250 mM NaCl, 50 mM MgCl_2_) were added to the suspension in order to obtain an isotonic environment. The suspension was gently mixed and transferred in a 2 ml Eppendorf tube using a glass Pasteur pipette and the sample was centrifuged at 1,000 x g at 4°C for 5 min to separate cells, debris, and nuclei. The supernatant was referred as Cyto1. The pellet was resuspended in 900 µl of HypoB and the same procedure was repeated. The supernatant was referred as Cyto2. Cyto1 and Cyto2 fractions were pooled together and centrifuged at 1,000 x g at 4 °C for 5 min to separate debris and the supernatant was collected and transferred in a 2 ml tube. After centrifugation at 15,000 x g at 4 °C for 2 min, the pellet (lysosomal fraction), referred as L, was resuspended in 200 µl of Suspension Buffer (10 mM HEPES, pH 7.4, 150 mM NaCl supplemented with phosphatase/protease inhibitor). The supernatant was transferred in a 2 ml Eppendorf tube and centrifuged at 17,000 x g at 4 °C for 20 min and the supernatant (cytosolic fraction) was marked as C. For both mice and HepG2-derived samples, 50 µl of L, C or EL fractions were dispensed in a 0.250 ml Eppendorf tube, supplemented with 5 µl 100 mM glycine (10 mM final concentration), or vehicle (water) and sealed with parafilm. Samples were incubated at 37 °C, for 1 h in an orbital shaker (450 RPM) and, after the incubation, the samples received 50 µl 1 M NaOH (500 mM final concentration; dispensed with an insulin syringe by punching the lid of the Eppendorf, without opening the tube).

Lysosome disruption was achieved by exposing L fraction to 5 cycles of 30 s pulse followed by 30 s pause on ice (using an ultrasonic bath sonicator), followed by 3 cycles of freeze (10 min at -20 °C) and thaw (3 min at 37 °C). Lysosomal disruption was confirmed by electron microscopy as described**^60^**. Briefly, isolated lysosomes (intact or disrupted) were fixed in 2.5% glutaraldehyde (Polysciences Inc, Glutaraldehyde EM Grade 25%, N° 01909-100) in PBS for at least 1 hour at room temperature, before centrifuged in a 1.5 ml Eppendorf tube in an Eppendorf centrifuge 5430R at 14000 rpm (rotor: FA-45-30-11) for 10 minutes at room temperature to get a visible pellet. After discarding the supernatant, the pellet was resuspended and post-fixed in 2% osmium tetroxide in H_2_O for 60 minutes at 4°C. After centrifugation (same conditions as described above), the pellet was embedded into epon (Sigma-Aldrich, Epoxy embedding medium, N°45345-250ML-F, Epoxy embedding medium hardener DDSA, N°45346-250ML-F, Epoxy embedding medium hardener MNA, N°45347-250ML-F, Epoxy embedding medium accelerator DMP30, N°45348-250ML-F). The samples were further processed for thin sections (90nm thickness), before staining with lead citrate and uranyl acetate (Fluka, Lead(II) nitrate, N°15334)+(Merck, Tri-sodium citrate dihydrate, N°1.06430). The samples were imaged with a CM100 transmission electron microscope (Philips, Eindhoven, The Netherlands). The samples with disrupted lysosomes contained substantially decreased vesicular structures with discontinued, damaged membrane with less or no luminal content, whereas all vesicles in the intact lysosome preparation had intact membranes with plenty of luminal content. Cyanide detection was performed with electrochemical and LC-MS/MS methods (see below). Remaining lysosomal samples were lysed with Cell-Lytic^TM^ M buffer supplemented with 1% phosphatase/protease inhibitor cocktail Halt, 1861281 and stored at -20 °C for further western blotting analysis.

*HCN generation by MPO or PXDN isolated enzymes.* MPO (200 ng/well) or PXDN (500 ng/well) cyanogenic activity assay was performed in a 50 mM phosphate-citrate buffer, pH 4.5, or PBS buffer, pH 7.4, containing 1 mM glycine, 1 mM H_2_O_2_ and 150 mM NaCl, in a final volume of 50 µl. Assay mixture was incubated for 30 min at 37 °C and the reaction was stopped by the addition of 50 µl of 1 M NaOH. Cyanide production was measured with the electrochemical method (see below).

*Non-enzymatic HCN generation.* Equimolar concentrations of hypochlorous acid (HOCl) and glycine (10 mM) or various proteinogenic amino acids, in 50 mM phosphate-citrate buffer, pH 3.5-5.0, were incubated in 0.25 ml Eppendorf tubes, in a final volume of 50 µl and sealed with rubber cap. Assay mixture was incubated for 10 min at RT and the reaction was stopped by the addition of 50 µl of 1 M NaOH with a Hamilton syringe (without removing the cap). Cyanide production was measured with the electrochemical method (see below).

*Electrochemical method for HCN detection (as CN^-^):* HCN levels were measured using a CN^-^ selective electrode (Lazar Research Labs, Inc., LIS-146CNCM-XS micro ion)**^9,61–63^**. Prior to measuring, all samples were prepared by diluting 1:1 (v/v) in 1 M NaOH (0.5 M, final concentration) and incubated at RT for 30 min, thus inducing the partition of HCN to CN^-^. The CN^-^ electrode was fully soaked within the sample and voltage (mV) was acquired until the signal was stabilized. The value of the blank sample was subtracted from all measurements. CN^-^ concentration was calculated against a standard curve; absolute cyanide concentrations were typically in the range of 1-20 µM. Data were normalized to total mg of protein obtained using BCA method (Thermo Scientific™, 23225) for cells and lysosomes, or total wet weight for mouse tissues and expressed as cyanide generation rate (nmoles cyanide/mg protein·h for cells and lysosomes or nmoles cyanide/mg tissue·h for wet tissues).

*HCN measurement using the Cyanalyzer LC-MS/MS method:* All solvents used were LC/MS grade. All reagents used were analytical standard grade. Potassium cyanide (KCN, CAS 151-50-8, CAT AAL1327336), sodium hydroxide (NaOH, CAS 1310-73-2, CAT S25548C), sulfuric acid (H_2_SO_4_, CAS 7664-93-9, CAT A300-212), ammonium formate (CAS 540-69-2, CAT A11550), potassium dihydrogen phosphate (KH_2_PO_4_, CAS 7778-77-0, CAT P386-500), and dibasic potassium phosphate (K_2_HPO_4_, CAS 7758-11-4, CAT P288-500) were purchased from Fisher Scientific (Hanover Park, IL, USA). Naphthalene-2,3-dicarboxaldehyde (NDA, CAS 7149-49-7, CAT A5594) was purchased from TCI America (Portland, OR, USA). 2-Aminoethane sulfonic acid (Taurine, CAS 107-35-7, CAT A12403.36) and sodium metaborate tetrahydrate (NaBO_2_·4H_2_O, CAS 10555-76-7, CAT A11803.36) were purchased from Alfa Aesar (Ward Hill, MA, USA). Purified water was obtained from a water PRO PS polisher (Labconco, Kansas City, KS, USA) at a resistivity of 18.2 MΩ-cm. Phosphate borate buffer (50 mM; pH 8.5) and NaOH (10 mM) were prepared in deionized water and transferred into plastic containers for bench-top storage. An NDA stock solution (2 mM) was prepared in 50 mM phosphate borate buffer and 40% methanol and stored in an amber vial at room temperature. Taurine (100 mM) solution was prepared in 50 mM phosphate-borate buffer and stored at room temperature. H_2_SO_4_ (2 M) was prepared in deionized water and 50% ethanol and stored at room temperature. The calibration standards were prepared from an aqueous CN stock solution (10 mM). All the calibration standards for CN (0.5, 1.0, 2.0, 5.0, 10, 20, 50, 100, and 200 μM) were prepared in PBS buffer (in case of LH samples) or whole blood (in case of blood samples). All calibrators were prepared in triplicate.

For the measurement of HCN generation from liver homogenates, glycine (7.5 µl of 200 mM aqueous solution) or its vehicle was added to 142.5 µl LH sample in a 0.5 ml screw cap vial. The vial was capped, vortexed, and incubated at 37 °C for 20 h in a benchtop Fisher Scientific Isotemp incubator. After incubation, 25 μl was removed from the vial and used for the analysis. Quantification of HCN proceeded via active microdiffusion, chemical modification of HCN using NDA and taurine, and LC-MS/MS analysis of the CN-NDA-taurine compound as described**^10,36,64,65^**. Briefly, NDA (2 mM), taurine (100 mM), and NaOH (10 mM), 200 µl each, were added to the reagent chamber of a two-chamber sample preparation cartridge which allows active airflow from the sample chamber to the reagent chamber. LH sample or mouse blood (25 µl) was placed in the sample chamber and diluted with 80 µl of deionized water. H_2_SO_4_ (200 µl of a 2 M aqueous solution) was added to the sample chamber to ensure HCN in the LH or mouse blood was in the gaseous (HCN) form. The sample and reagent chambers were immediately capped. Carrier gas (i.e., room air at 200 ml/min) was pumped through the sample chamber into the capture chamber to transfer HCN gas to the capture solution. In the capture chamber, the NDA and taurine reacted with HCN to form a CN-NDA-taurine complex. An aliquot (250 µl) of the CN capture chamber solution was filtered with a 0.22-μm polytetrafluoroethylene (PTFE) into a 300-μl glass insert placed in a 2 ml HPLC vial for subsequent HPLC–MS/MS analysis. Prepared samples were analyzed using a Shimadzu HPLC (LC20AD, Shimadzu Corp., Kyoto, Japan). Separation was achieved by reversed-phase chromatography using a ZORBAX RRHT Eclipse Plus C18 column (100 × 3.0 mm, 1.8 μm, 95 Å). Each chromatographic analysis was carried out using 10 mM aqueous ammonium formate (mobile phase A) and 10 mM ammonium formate in methanol (mobile phase B). An aliquot (10 μl, injection volume) of the prepared sample was separated with gradient elution at a flow rate of 0.3 ml/min and column temperature of 40 °C as follows: the column was initially equilibrated with 50% mobile phase B, linearly increased to 100% mobile phase B over 3 min, maintained at 100% B for 1 min, and then decreased linearly back to 50% mobile phase B over 1 min, where the mobile phase composition was held constant for 2 min to re-equilibrate the column between samples. A Sciex 5500 Q-trap MS/MS (Applied Biosystems, Foster City, CA, USA) with multiple reaction monitoring (MRM) was used to detect the CN-NDA-taurine complex using electrospray ionization (ESI) operated in negative ion mode. Nitrogen (20 psi) was used as the curtain gas. The ion source was operated at −4,500 V, a temperature of 500 °C, and a pressure of 10 psi and 0 psi for nebulizer and drying gases, respectively. The m/z ratio of 298.6 corresponds to the molecular ion of the CN-NDA-taurine complex. The transitions 298.6→190.7 and 298.6→80.9 were used as the quantification and identification transitions, respectively. The collision cell was operated with a medium collision gas pressure and an entrance potential of −10 V, a declustering potential of −70.0 V, a collision energy of −25.0 V, a collision exit potential of −19.0 V, and a dwell time of 100 ms. Blank values were subtracted from all measured values. HCN concentration was calculated against a standard curve; absolute concentrations measured were typically in the range of 30-300 nM. Data were normalized to total mg of protein obtained using BCA method (Thermo Scientific™, 23225) for lysosomes, or total wet weight for mouse tissues.

*HCN measurement using a monocyano-cobinamide (MCC) based spectrophotometric method:* HCN was measured using a spectrophotometric method which exploits absorption changes that occur on conversion of monocyano-cobinamide (MCC) to dicyano-cobinamide at 366 nm**^66^**. The reactions were carried out in a modified 96-deep well-plate, with a communication bridge between adjacent wells and covered by a silicon lid. This creates a closed system enabling gas exchange between two communicating wells. The polypropylene 2-ml deep square 96-well plates (JTBaker/Avantor, 43001-0020) were manually modified using Dremel tool to connect two horizontally adjacent wells by a small hole cca. 2-4 mm in diameter in a dividing wall around 3 mm from the top edge. Wells on the left served to carry out the activity assay, while wells on the right contained 200 µl of 100 µM monocyanocobinamide (MCC) in 100 mM NaOH. The plate was air-tight sealed using ethylene-vinyl acetate capmat for square 96-well plate (JTBaker/Avantor, 43001-0022). To stop the reaction and for a complete volatilization of HCN from enzymatic reactions, 500 µl of 0.5 M H_2_SO_4_ was injected into the wells on the left using a syringe equipped with a 26G needle via self-sealing capmat and plate was incubated at 37 °C overnight. Next morning, the capmat was carefully removed, MCC solution from wells on the right was transferred into a clear polystyrene 96-well plate and read at 366 nm using Spectramax M5 plate reader (Molecular Devices). Blank values were subtracted from all measured values. HCN concentrations were calculated against a standard curve, the absolute concentrations measured were typically in the range of 30-100 nM. Data were normalized to total wet weight for mouse tissues.

### Confocal microscopy

*Live cell imaging*: HepG2 cells were seeded on poly-L-ornithine coated glass-bottom microscopy chamber at a density 300,000 cells/well and incubated overnight. Human primary hepatocytes were seeded (in plating medium) on collagen coated glass-bottom microscopy chamber at a density 150,000 cells/well and incubated for 6 h. Human primary hepatocytes (in maintenance medium) and HepG2 cells were treated with 10 mM glycine for 24 h and 10 µM THC for 3 h. Before the experiment cells were incubated for 1 hour with 10 µM of either the fluorescent HCN probe CSP, a spiropyrane derivatives of cyanobiphenyl**^14^** which undergoes a turn-on fluorescence in the presence of HCN (Ex 405 nm / Em 495 nm) or the fluorescent HCN/HOCl probe Chemosensor P (2-amino-3-((5-(benzothiazol-2-yl)-2-hydroxybenzylidene)amino) maleonitrile)**^15^** which undergoes a sharp turn-on fluorescence in the presence of HCN (Ex 405 nm / Em 584-620 nm) or in the presence of HOCl (Ex 405 nm / Em 450-550 nm). For co-localization experiments cells were also incubated together with cell-permeant dyes (50 nM LysoTrackerGreen, Thermo Fisher Scientific, L7526; 10 µM calcein, AM, Thermo Fisher Scientific, C3100; 1 µM CellMask™ Green Actin Tracking Stain, Thermo Fisher Scientific, A57247 or 200 nM MitoTracker Deep Red FM, Thermo Fisher Scientific, M22426) for 30 min at 37 °C and 5% CO_2_. At the end of the incubation, cells were washed 3-times and visualized using confocal microscope Leica SP5 or Leica 8 Stellaris Falcon using a 40 × oil-immersion APO Plan objective. Following excitation and emission spectra were used: LysoTrackerGreen (Ex 488 nm / Em 517 nm), calcein, AM (Ex 488 nm / Em 517 nm), CellMask™ Plasma Membrane Stain (Ex 488 nm / Em 535 nm) MitoTrackerDeepRed (Ex 644 nm / Em 665 nm). The parameters of acquisition: image format of 1,024 × 1,024 pixels, 200 Hz scan speed. Cell fluorescence was measured using ImageJ using the corrected total cell fluorescence (CTCF) formula where CTCF is defined as Integrated Density-(area of selected cell x mean fluorescence of background reading).

*MPO localization and subcellular organelles:* HepG2 cells were seeded on poly-L-ornithine coated cover glass in 12-well plate at a density 100,000 cells/well and incubated with the appropriate medium until 40–60% confluence was reached. Cells were loaded with 500 nM ER tracker green for 30 min. Cells were then washed twice with pre-warmed 0.1 M TBS (Tris-buffered saline), fixed with pre-warmed 4% paraformaldehyde solution for 2 min at 37 °C and incubated in blocking buffer (0.1 M TBS containing 10% donkey serum) for 60 min at room temperature. Primary anti-MPO antibody (rabbit, dilution 1:10,000, Sigma Aldrich, HPA021147) was added and incubated overnight at 4 °C. The following day, cells were washed three times with TBS and incubated with appropriate secondary antibody goat anti-rabbit IgG (H+L) Highly Cross-Adsorbed Secondary Antibody Alexa Fluor Plus 647 (1:1,000 dilution), were added for 1 h at room temperature. DAPI (Molecular Probes, 5 µg/ml, Thermo Fisher Scientific, D1306) was added for the last 5 minutes. Cells were then washed three times and coverslipped with Prolong Gold antifade reagent (Thermo Fisher Scientific, P36930) and visualized using Leica SP5 or Leica 8 STELLARIS Falcon at 63x magnification. ERtracker was visualized at Ex 504 nm / Em 511 nm and MPO at Ex 647 nm / Em 665 nm.

For experiments with MitoTracker Deep Red, cells were loaded with 200 nM MitoTracker Deep Red FM and fixed with 4% PFA for 15 min at 37 °C and incubated in blocking buffer (0.1 M TBS containing 10% donkey serum) for 60 min at room temperature. Primary antibody anti-MPO antibody (rabbit, dilution 1:10,000, Sigma-Aldrich, HPA021147) was added and incubated overnight at 4 °C. The following day, cells were washed three times with TBS and incubated with appropriate secondary antibody goat anti-rabbit IgG (H+L) Highly Cross-Adsorbed Secondary Antibody Alexa Fluor Plus 568 (1:1,000 dilution, anti-mouse Alexa Fluor™ 568, Thermo Fisher Scientific, A-11004), were added for 1 h at room temperature. DAPI (Molecular Probes, 5 µg/ml, Thermo Fisher Scientific, D1306) was added for the last 5 min. Cells were then washed three times and coverslipped with Prolong Gold antifade reagent (Thermo Fisher Scientific, P36930) and visualized using Leica 8 STELLARIS Falcon at 63x magnification. MitoTracker was visualized at Ex647/Em 665 nm and MPO at Ex 568 nm.

### Proteomics and detection of S-cyanylation by LC-MS/MS

*Mouse liver tissue treated with 10 mM glycine or vehicle:* Mouse liver samples (50 mg each) were obtained frozen at -80 °C. Livers were cut in half using a scalpel and forceps and weighed on a scale. Tissues were added to a Potter homogenizer with 2 ml of PBS pH 7.4 (Dulbecco′s Phosphate Buffered Saline, Sigma-Aldrich, D8537) and homogenized until completely broken and mixed with PBS. 1 ml of the sample was transferred to a 5 ml Eppendorf to which 1 ml of 20 mM glycine (either standard “light” C^12^N^14^glycine, or “heavy” C^13^N^15^-glycine, both from Sigma/Aldrich) was added. The other half of the sample was also transferred and 1 ml of PBS was added. In each sample was added protease inhibitor to volume of 1%. Samples were placed in a 6 well plate and put into the incubator (37 °C, 5% CO_2_) for 8 hours. Samples were transferred to 5 ml Eppendorf tubes and to each was added 1 ml of 2X HEN (100 mM HEPES, 2 mM EDTA, 0.2 mM neocuproine, 2% IGEPAL, 4% SDS, pH 7.4) with 2% protease inhibitor and 40 mM iodoacetamide. Samples were lysed using a handheld homogenizer and precipitated with methanol/chloroform method. Pellets were left to dry overnight. Dry samples were resuspended using 50 µl 50 mM ABC, 6 M guanidine HCl. 20 µg of protein was diluted 20 times with 50 mM ABC (no guanidine) and trypsin was added with a 1:20 trypsin to sample ratio (Promega, V5117). Protein were digested overnight at 37 °C. The desalting was performed on Oasis HLB 1 cc Vac Cartridges, 30 mg sorbent (Waters, WAT094225). Columns were washed with the elution solvent 60% acetonitrile in 0.1% trifluoroacetic acid (0.1% TFA) and then twice with 0.1% TFA. Samples were then loaded on the column by gravity flow. Cartridges were washed once with 1 ml 0.1% TFA, followed by two times gravity flow elution with the elution solvent. Desalted samples were evaporated under vacuum until dryness. Dry samples were resuspended in 40 µl 0.1% TFA and the quality control was performed using an Ultimate 3000 Nano Ultra High-Pressure Chromatography (UPLC) system with a PepSwift Monolithic® Trap 200 µm * 5 mm (Thermo Fisher Scientific). Peptides were analyzed on high-resolution LC-MS/MS using an Ultimate 3000 Nano Ultra High-Pressure Chromatography (UPLC) system (Thermo Fisher Scientific) coupled to a timsTOF Pro (Bruker) equipped with a CaptiveSpray source. Peptide separation was carried out with an Acclaim™ PepMap™ 100 C18 column (Thermo Fisher Scientific) using a 150 min linear gradient from 3 to 42% of B (84% acetonitrile, 0.1% formic acid) at a flow rate of 250 nl/min. Data were evaluated with PEAKS ONLINE software using 20 ppm for precursor mass tolerance, 0.5 Da for fragment mass tolerance, specific tryptic digest, and a maximum of 3 missed cleavages. Carbamidomethylation (+57.021464 Da) on C, N-term acetylation (+42.010565 Da), methionine oxidation (+15.994915 Da), and on cyanylated peptides only Cyano PTM (+24.995249 Da) were added as variable modifications, PSM and proteins were filtered at FDR 1 %. Data were normalized based on the total ion count (TIC).

*HepG2 cells treated with 10 mM glycine and serine/glycine free medium:* HepG2 cells (500,000/well) were seeded in 6-well plates and incubated overnight at 37 °C and 5% CO_2_. The day after cells were treated with 10 mM glycine, -Ser/Gly medium and control and further incubated for 24 h. After the incubation, cells were washed in ice-cold PBS and lysed in lysis buffer (CelLytic^TM^ M, Sigma-Aldrich, C2978) supplemented with 1% protease inhibitor and 20 mM iodoacetamide. Samples were incubated at 37 °C for 1 h, 450 RPM. Samples were centrifuged for 10 min at max rcf (20,000 rcf) at 4 °C, the supernatant was taken and proteins were precipitated with methanol/chloroform method. Proteins were pelleted by methanol/chloroform precipitation protocol. Samples were resuspended in 50 mM ABC 6 M guanidine HCl to a concentration of 6 mg/ml. Their protein concentration was determined using DC protein assay (Lowry method). 20 µg of protein was diluted 20 times with 50 mM ABC (no guanidine) and trypsin was added with a 1:20 trypsin to sample ratio (Promega, V5117). Protein were digested overnight at 37 °C. The desalting was performed on Oasis HLB 1 cc Vac Cartridges, 30 mg sorbent (Waters, WAT094225). Columns were washed with the elution solvent 60% acetonitrile in 0.1% trifluoroacetic acid (0.1% TFA) and then twice with 0.1% TFA. Samples were then loaded on the column by gravity flow. Cartridges were washed once with 1 ml 0.1% TFA, followed by two times gravity flow elution with the elution solvent. Desalted samples were evaporated under vacuum until dryness. Dry samples were resuspended in 40 µl 0.1% TFA and the quality control was performed using an Ultimate 3000 Nano Ultra High-Pressure Chromatography (UPLC) system with a PepSwift Monolithic® Trap 200 µm * 5 mm (Thermo Fisher Scientific). Peptides were analyzed on high-resolution LC-MS/MS using an Ultimate 3000 Nano Ultra High-Pressure Chromatography (UPLC) system (Thermo Fisher Scientific) coupled to a timsTOF Pro (Bruker) equipped with a CaptiveSpray source. Peptide separation was carried out with an Acclaim™ PepMap™ 100 C18 column (Thermo Fisher Scientific) using a 150 min linear gradient from 3 to 42% of B (84% acetonitrile, 0.1% formic acid) at a flow rate of 250 nl/min. Data were evaluated with PEAKS ONLINE software using 20 ppm for precursor mass tolerance, 0.5 Da for fragment mass tolerance, specific tryptic digest, and a maximum of 3 missed cleavages. Carbamidomethylation (+57.021464 Da) on C, N-term acetylation (+42.010565 Da), methionine oxidation (+15.994915 Da), and on cyanylated peptides only Cyano PTM (+24.995249 Da) were added as variable modifications, PSM and proteins were filtered at FDR 1 %. Data were normalized based on the total ion count (TIC).

*HepG2 cells treated with HCN-releasing agents:* HepG2 cells (500,000/well) were seeded in 6-well plates and incubated overnight at 37 °C and 5% CO_2_. The day after cells were treated with 30 µM amygdalin, 100 µM linamarin, 100 µM mandelonitrile, 1 µM KCN or 10 nM KCN and further incubated for 24 h. After the incubation, cells were washed in ice-cold PBS and lysed in HEN lysis buffer (50 mM HEPES, 1 mM EDTA, 0.1 mM neocuproine, 1% IGEPAL, 2% SDS, pH 7.4) supplemented with 1% protease inhibitor and 20 mM iodoacetamide using syringes. Samples were then left to incubate in the dark at 37 °C for 1.5 h. Samples were centrifuged for 15 min at max rcf (20,000 rcf) at 4 °C, the supernatant was taken and proteins were precipitated with methanol/chloroform method. Pellets were resuspended in 120 µl of 50 mM ABC 1 M guanidine HCl. 20 µg of protein was diluted 10 times with 50 mM ABC (no Guanidine) and trypsin was added with a 1:20 trypsin to sample ratio (Promega, V5117). Protein were digested overnight at 37 °C. The desalting was performed on Oasis HLB 1 cc Vac Cartridges, 30 mg sorbent (Waters, WAT094225). Columns were washed with the elution solvent 60% acetonitrile in 0.1% Trifluoroacetic acid (0.1% TFA) and then twice with 0.1% TFA. Samples were then loaded on the column by gravity flow. Cartridges were washed once with 1 ml 0.1% TFA, followed by two times gravity flow elution with the elution solvent. Desalted samples were evaporated under vacuum until dryness. Dry samples were resuspended in 40 µl 0.1% TFA and the quality control was performed using an Ultimate 3000 Nano Ultra High-Pressure Chromatography (UPLC) system with a PepSwift Monolithic® Trap 200 µm * 5 mm (Thermo Fisher Scientific). Peptides were analyzed on high-resolution LC-MS/MS using an Ultimate 3000 Nano Ultra High-Pressure Chromatography (UPLC) system (Thermo Fisher Scientific) coupled to a timsTOF Pro (Bruker) equipped with a CaptiveSpray source. Peptide separation was carried out with an Acclaim™ PepMap™ 100 C18 column (Thermo Fisher Scientific) using a 120 min linear gradient from 3 to 30% of B (84% acetonitrile, 0.1% formic acid) at a flow rate of 250 nl/min. Data were evaluated with PEAKS ONLINE software using 20 ppm for precursor mass tolerance, 0.5 Da for fragment mass tolerance, specific tryptic digest, and a maximum of 3 missed cleavages. Carbamidomethylation (+57.021464 Da) on C, N-term acetylation (+42.010565 Da), methionine oxidation (+15.994915 Da), and on cyanylated peptides only Cyano PTM (+24.995249 Da) were added as variable modifications, PSM and proteins were filtered at FDR 1 %. Data were normalized based on the total ion count (TIC).

*Cyanylation of GAPDH or GPDH:* GAPDH (0.14 mg/ml in PBS buffer) was treated with 10 µM KCN, 10 µM H_2_O_2_ and both KCN and H_2_O_2_ for 30 min at RT. GPDH (0.14 mg/ml in PBS buffer) was treated with 30 µM KCN, 30 µM H_2_O_2_ and both KCN and H_2_O_2_ for 30 min at RT. 1 µg of protein was diluted 5 times with 50 mM ABC and trypsin was added with a 1:20 trypsin to sample ratio (Promega, V5117). Protein were digested 24 h at 37 °C. The desalting was performed on Oasis HLB 1 cc Vac Cartridges, 30 mg sorbent (Waters, WAT094225). Columns were washed with the elution solvent 60% acetonitrile in 0.1% trifluoroacetic acid (0.1% TFA) and then twice with 0.1% TFA. Samples were then loaded on the column by gravity flow. Cartridges were washed once with 1 ml 0.1% TFA, followed by two times gravity flow elution with the elution solvent. Desalted samples were evaporated under vacuum until dryness. Dry samples were resuspended in 4 µl 0.1% TFA and the quality control was performed using an Ultimate 3000 Nano Ultra High-Pressure Chromatography (UPLC) system with a PepSwift Monolithic® Trap 200 µm * 5 mm (Thermo Fisher Scientific). Peptides were analyzed on high-resolution LC-MS/MS using an Ultimate 3000 Nano Ultra High-Pressure Chromatography (UPLC) system (Thermo Fisher Scientific) coupled to a timsTOF Pro (Bruker) equipped with a CaptiveSpray source. Peptide separation was carried out with an Acclaim™ PepMap™ 100 C18 column (Thermo Fisher Scientific) using a 120 min linear gradient from 3 to 35% of B (84% Acetonitrile, 0.1% Formic Acid) at a flow rate of 250 nl/min. Data were evaluated with PEAKS ONLINE software using 20 ppm for precursor mass tolerance, 0.5 Da for fragment mass tolerance, specific tryptic digest, and a maximum of 3 missed cleavages. Carbamidomethylation (+57.021464 Da) on C, N-term acetylation (+42.010565 Da), methionine oxidation (+15.994915 Da), and on cyanylated peptides only Cyano PTM (+24.995249 Da) were added as variable modifications, PSM and proteins were filtered at FDR 1%. Data were normalized based on the total ion count (TIC).

In the mouse liver we have detected 248 quantifiable peptides belonging to 213 proteins. Among those, 71 peptides (67 proteins) are increasing upon glycine treatment. In HepG2 cells, we have identified and quantified 192 peptides corresponding to 170 proteins. Among them, 64 peptides (60 proteins) were increases upon glycine treatment. On the contrary, in response to glycine-serine free media incubation, we have detected 206 peptides (180 proteins) for which 57 (or 48 proteins) were decreasing. 192 peptides corresponding to 168 proteins were quantifiable. 42 peptides (42 proteins) were increasing with amygdalin treatment. 65 peptides (64 proteins) in linamarin treatment and 77 peptides (75 proteins) in mandelonitrile were also found increasing. Similar number of increasing peptides were found after treatment with 1 µM and 10 nM KCN, 75 peptides (70 proteins) and 70 peptides (67 proteins) respectively. Among all those changes only 12 peptides /protein were found in common. Although there are no methods for selective labeling and enrichment of cyanylated proteins – and consequently, no enrichment was applied in this study – cyanylated proteins could be readily detected. The numbers of cyanylated proteins we report likely underestimate the total number of proteins in the cells and tissues investigated.

### Detection of protein S-cyanylation by SDS-PAGE

This method is based on the principle that chemical cleavage of polypeptide’s backbone occurs after S-cyanylation of target cysteine residues under alkaline and denaturating conditions and has been previously employed to detect S-cyanylation of various plasma proteins**^67–69^**. Briefly, in PCR tubes isolated GAPDH or GPDH 0.14 mg/ml in PBS buffer was pre-treated with 1 mM KCN or 0.325 mM H_2_O_2_ (used to induce oxidation of cysteine residues) or 0.325 mM diamide (used to induce oxidation of cysteine residues specifically to disulfide) at RT for 30 min under constant agitation (750 RPM). Afterwards, samples were treated with 1 mM KCN or 0.325 mM H_2_O_2_ or 0.325 mM diamide and further incubated at RT for 30 min under constant agitation (750 RPM). The reaction mixture was alkalinized by adding NaOH, yielding a final concentration of 0.12 M, and resuspended in an isovolume of 2x Lämmli buffer supplemented with 5% β-mercaptoethanol, thus inducing protein denaturation. Samples were run against a polyacrylamide gel electrophoresis (SDS-PAGE), followed by Coomassie staining.

### Hypoxia and hypoxia-reoxygenation experiments

HepG2 cells were seeded in 96-well plate at 20,000 cells/well or in a 6-well plate at 500,000 cells/well and incubated at 37 °C and 5% CO_2_ overnight. To monitor the effect of glycine and HCN scavengers, the day after seeding medium was replaced with -Ser/Gly medium or with -Ser/Gly medium supplemented with 0.4 mM glycine or with -Ser/Gly medium supplemented with 10 mM glycine. Cells were moved to hypoxic chamber (InvivO2 400, Baker Ruskinn, Bridgend, UK) at 0.2% O_2_ and 5% CO_2_ and subsequently incubated for 48h. For the last 3 h, a group of cells were treated with HCN scavengers (THC, 10 µM or CoE, 10 µM); control cells received vehicle. To monitor the effect of HCN and HCN releasers, the day after seeding medium was replaced with medium supplemented with 10 nM KCN or with HCN releasers amygdalin (30 µM), linamarin (100 µM) or mandelonitrile (100 µM) and cells were incubated for 48 h under hypoxic conditions (0.2% O_2_). In all cases, reoxygenation was also tested by incubating cells under normoxic conditions (5% CO_2_) for an additional 24 h period.

*Cell viability assessment:* After treatments and incubation, 50 µl of each well’s supernatant was transferred to another plate to test lactate dehydrogenase (LDH) activity, an indicator of cell necrosis. The LDH assay was performed using the Pierce LDH Cytotoxicity Detection Kit Plus (Roche, 04744926001). Briefly, 50 µl/well of LDH reaction mixture were added to the supernatants. The plate was incubated for 30 min at RT protected from light and the reaction was stopped with 25 µl/well of Stop Solution. The plate was shaken in an orbital way for 60 s before measuring the absorbance at 490 nm and 680 nm (background) with an Infinite 200 Pro reader (Tecan, Männedorf, Switzerland).

*Sample preparation for Western blotting:* After treatments and incubation cells seeded in 6-well plate were washed with 500 µl/well PBS and, to preserve the stability of HIF-1α, cells were lysed directly with 200 µl LDS Sample Buffer (1X) (Invitrogen, 2134101) supplemented with Reducing Agent (1X) (Invitrogen, 199821). Samples were collected by scraping the wells and sonicated for 10 min (10 cycles of 30 s sonication/30 s stop at RT). Samples were incubated for 10 min at 95 °C and were immediately loaded in 4%–12% Bis-Tris Plus Gels (Invitrogen, Thermo Scientific) and ran at constant 120 V (see below).

### Western blot analysis

Western blot analysis was performed according to standard protocols. Briefly, 20 μg of total protein (from whole cell extracts or isolated lysosomes) was separated with a 4%–12% Bis-Tris Plus Gels (Invitrogen, Thermo Scientific) and then transferred onto PVDF (polyvinylidene difluoride) membrane by dry transfer using the iBlot™ 2 Device and Transfer Stacks (Invitrogen). After blocking with 5% Milk in TBS-Tween20 (TBS-T) for 1 h, membranes were incubated with specific antibodies overnight at 4 °C. The following antibodies were used: anti-HIF-1α mouse monoclonal antibody (BD Transduction Laboratories, 610958, 1:1,000), anti-LAMP1 antibody rabbit monoclonal (Abcam, ab225762, 1:1,000), anti-GAPDH rabbit polyclonal (Sigma-Aldrich, ABS16, 1:1,000), anti-myeloperoxidase rabbit monoclonal (E1E7I) (Cell Signaling Technology, 14569, 1:1,000), anti-catalase (D4P7B) (Cell Signaling Technology, 12980, 1:1,000), anti-myc tag (9B11) (Cell Signaling Technology, 2278, 1:1,000), anti-β-actin mouse monoclonal (AC-15) (Sigma-Aldrich, A1978, 1:1,000), anti-PXDN rabbit polyclonal (Sigma-Aldrich, ABS1675, 1:1,000), anti-TST rabbit polyclonal (Abcam, ab231248, 1:1,000). Rabbit polyclonal anti-MGST1, anti-GSTA1, anti-GSTA2, anti-PRDX3 and anti-PRDX6 were purchased from GeneTex (Irvine, CA, USA). After incubation, the membranes were briefly washed 3 times with TBS and incubated for 1 h in room temperature with secondary antibodies anti-rabbit IgG or anti-mouse IgG, HRP-linked antibody (Cell Signaling, 7076) diluted 1:3,000 in TBS-T/5% Milk. The membranes were washed twice with TBS-T and once with TBS and the detection was performed with Amersham ECL™ Prime Western Blotting Detection Reagent (GE Healthcare). Chemiluminescence was detected with the Azure Imaging System 300 (Azure Biosystems, Dublin, CA, USA).

### Bioenergetic analysis in HepG2 cells

The Seahorse XFe24 flux analyzer (Agilent Technologies, Santa Clara, CA, USA) was used**^70^** to estimate cellular bioenergetics of HepG2 cells or dermal fibroblasts from Detroit551, G;00880, GM00747, GM10360. HepG2 cells were seeded at a density of 20,000 cells/well on Agilent Seahorse XF24 cell culture microplates. The day after, medium was replaced with fresh medium supplemented with 10 mM glycine or -Ser/Gly medium and further incubated for 24 h. To reduce HCN levels, cells were treated with HCN scavengers THC or CoE (10 µM) and incubated at 37 °C and 5% CO_2_ for 3 h. NKH cells or control dermal fibroblasts were incubated with vehicle or hydroxychloroquine (30 µM) for 72h followed by seeding at a density of 5,000 cells/well on Agilent Seahorse XF24 cell culture microplates for bioenergetic measurement.

*Mito Stress Test:* For analysis of mitochondrial respiration, the Cell Mito Stress assay kit (Agilent, 103015-100) was used. Cells were washed twice with DMEM (pH 7.4) supplemented with L-glutamine (2 mM, Corning, 25-015-CL), sodium pyruvate (1 mM, Cytiva, SH30239.01) and glucose (10 mM, Sigma-Aldrich, G8644). The microplate was then incubated in a CO_2_-free incubator at 37 °C for 1 h, to allow temperature and pH equilibration, as recommended by the manufacturer. The assay consisted in two measurements of basal values of oxygen consumption rate (OCR), followed by the injection of 1 μM oligomycin, used to evaluate ATP generation rate. Subsequently, the mitochondrial oxidative phosphorylation uncoupler FCCP (0.8 μM for HepG2, 2 μM for dermal fibroblasts), was employed to estimate maximal mitochondrial respiratory capacity. Eventually, 0.5 μM of rotenone and antimycin A were injected to inhibit the electron flux through the complex I and III, respectively, aiming to detect the extra-mitochondrial OCR.

*Long Chain Fatty Acid Oxidation Stress Test:* For the analysis of the fatty acid oxidation pathway, the Long Chain Fatty Acids Oxidation Kit (Agilent, 103672-100) was used. Cells were washed twice with DMEM (pH 7.4) supplemented with glucose (2 mM, Sigma-Aldrich) and L-carnitine (0.5 mM). After 1 h incubation at 37 °C in CO_2_-free, just prior starting the assay, each well received 87.5 µl of 1mM XF palmitate-BSA FAO substrate (Agilent, 102720-100) and etomoxir (4 µM, carnitine palmitoyl transferase-1 inhibitor) to reach a final volume of 500 µl/well. Measurements of basal values of oxygen consumption rate (OCR), were followed by injection of 1 μM oligomycin, followed by FCCP (0.8 μM). Subsequently, 0.5 μM of rotenone and antimycin A were injected to inhibit the electron flux through the complex I and III, respectively, aiming to detect the extra-mitochondrial OCR. Data were analyzed with Wave (v. 2.6; Agilent Technologies, Santa Clara, California, USA) and graphed with GraphPad Prism 8 (GraphPad Software Inc., San Diego, California, USA).

### Cell proliferation and viability

HepG2 cells were seeded in sterile transparent 96-well plate at 5,000 cells/well in 100 µl of complete culture medium and incubated over-night at 37 °C and 5% CO_2_. The day after, different treatments were added as indicated and cell proliferation was monitored for 72 h using the IncuCyte Live Cell Analysis device (20x objective) (Essen Bioscience, Hertfordshire, UK) as described**^71^**. Cell confluence was recorded every hour by phase contrast scanning for 72 h at 37 °C and 5% CO_2_ and calculated from the microscopy images. Data were analyzed using the IncuCyte® ZOOM v.2018A software (Essen Bioscience, Hertfordshire, UK).

Dermal fibroblasts were seeded at a density of 5,000 cells/well in sterile transparent 96-well plate and incubated for 72h. Every 24h, culturre medium was replaced with fresh medium containing vehicle (CTR) or 10 µM hydroxychloroquine (Hcq). End point cell proliferation was assayed with ELISA 5-bromo-2′ -deoxyuridine (BrdU) kit from Roche Diagnostics (Sigma– Aldrich). Briefly, cells were incubated with BrdU labelling solution for 2 h at 37 °C and 5% CO_2_. After removing the culture medium, DNA denaturation-fixation was performed, followed by incubation with the anti-BrdU antibody. Subsequently, the colorimetric substrate reaction was measured. Absorbance was measured at 450 nm with 690 nm as the reference wavelength using an Infinite M200 Pro plate reader (Tecan, Mannedof, Switzerland). The absorbance values directly correlate to the amount of DNA synthesized, and therefore is function of the number of proliferating cells in the respective microcultures. Concerning cell, viability, following cell treatment (as described above), dermal fibroblasts were further incubated for 1 h with 0.5 mg/ml MTT. The converted formazan dye was dissolved in 100 µl DMSO and the absorbance was measured at 570 nm and 690 nm (reference wavelength) with an Infinite 200 Pro plate reader (Tecan, Männedof, Switzerland).

### Measurement of intracellular glycine content

Glycine concentration in homogenates of HepG2 cells or fibroblasts (control and NKH) was measured by a fluorometric Glycine Assay Kit (Abcam, ab211100).

### GAPDH activity assay

The GAPDH Activity Assay Kit (Sigma-Aldrich, MAK277-1KT) has been used to measure the enzyme activity of glyceraldehyde-3-phosphate dehydrogenase enzyme after S-cyanylation. Prior to performing any experiment, GAPDH was incubated with 20 molar excess of DTT to favor reduced state of the enzyme. Excess of DTT was removed by employing a PD10 desalting column pre equilibrated in buffer Tris HCl 50 mM, pH 7.4. The reaction mixture, in the assay buffer provided with the kit, contained human GAPDH 0.55 µM and 10 µM H_2_O_2_, 10µM KCN or their combination, in a total assay volume of 50 µl. The 96-well plate was sealed with a transparent ELISA film and incubated for 30 min at RT. The enzymatic activity was triggered by the addition of a mixture containing substrates and developer. The activity was monitored at 450 nm, at 37 °C for 30 min using Infinite 200 Pro reader (Tecan, Männedorf, Switzerland).

### GPDH activity assay

Cytosolic glycerol-3-phosphate dehydrogenase (GPDH) (Merk, 10127752001) from rabbit muscle was used to test GPDH enzymatic activity after S-cyanylation. Prior to performing any experiment, GPDH was incubated with 20 molar excess of DTT to favor reduced state of the enzyme. Excess of DTT was removed by employing a PD10 desalting column pre equilibrated in buffer Tris HCl 50 mM, pH 7.4. The reaction mixture, in Tris HCl 50 mM pH 7.4 supplemented with EDTA 1mM, contained 0.05 µM GPDH, and 10 µM H_2_O_2_, 10µM KCN or a combination of them, in a total assay volume of 50 µl. The 96-well plate was sealed with a transparent ELISA film and incubated for 30 min at room temperature. The enzymatic activity was triggered by adding 500 µM NADH followed by 1.5 mM dihydroxyacetone phosphate. The oxidation of NADH to NAD^+^ was followed by monitoring the decrease in absorbance at 340 nm over time, at 37 °C using Infinite 200 Pro reader (Tecan, Männedorf, Switzerland).

### Metabolomics analysis

HepG2 cells were seeded in a sterile 10 ml dish at 2,000,000 cells/dish and incubated at 37 °C and 5% CO_2_. The day after, medium was replaced with the fresh medium supplemented with 10 mM glycine and further incubated for 24 h. Cells were washed twice with 10 ml of PBS and snap-frozen. Metabolites were quantified by LC-MS at the Metabolomics Unit of the University of Lausanne (Switzerland).

### RNA-seq

HepG2 cells were seeded in 6-well plate at 500,000 cells/well and incubated at 37 °C and 5% CO_2_. Next day, the medium was replaced with fresh medium supplemented with 10 mM glycine and further incubated for 24 h and harvested for analysis. For the comparison of TST silencing or TST overexpression, cells (wild-type control, TST-OE and shTST HepG2 cells) were seeded as above and incubated with normal cell culture medium containing 400 µM glycine for 24h and harvested for analysis. Cells were washed twice and RNA extraction and purification were performed using NucleoSpin^®^ RNA plus (Macherey-Nagel, 740984.250) according to the manufacturer’s instructions. RNA concentration and purity were measured with a nanodrop 2000 spectrophotometer (Thermo Scientific) by measuring the absorbance at 260/280 nm ratio. Samples were processed and analyzed using next generation sequencing (NGS) at Azenta Life Sciences (Griesheim, Germany). Fast gene set enrichment analysis (fGSEA) was performed on the complete (normalized) count data using the hallmark gene sets**^72^** using the GSEA_v.4.3.2 software (UC San Diego, La Jolla, CA, USA).

### Statistical analysis

If not otherwise stated, data are presented as mean values ± SEM of at least n=5 independent experiments where independent experiment is defined as an experiment performed on a different experimental day, as opposed to technical replicates. Differences among data are considered significant for p values < 0.05. Statistical analysis was performed using Student’s t-test to identify significant differences within the same group. Alternatively, two-way ANOVA followed by post-hoc Bonferroni’s multiple-comparison test was used to identify significant differences between two different groups.

## Acknowledgements

The authors thank Dr. Anita Marton for the editorial assistance. The authors also thank Mrs. Pauline Blanc for her excellent technical support for the electron microscopy work. We also thank the Transfusion CRS Fribourg (Fribourg, Switzerland) for the donation of whole blood leukoreduction filters.

## Authors’ contributions

Conceptualization: KZ, MP, CS

Methodology, Experimentation, Data Analysis: KZ, MP, LJ, AA, VM, KA, TM, SS, TMP, TV, JP, MK, SE, SM, DM, LF, SR, BS, TK, AKR, SC, GH.

Visualization: LJ, LF.

Funding acquisition: CS, MRF

Project administration: CS

Supervision: SC, CH, GH, AP, BAL, MRF, CS

Writing – original draft: KZ, MP, CS.

Writing – review & editing: KZ, MP, BAL, GRB, MRF, AP, DH, SC, CS.

## Data and materials availability

All data and materials are available from the corresponding author upon reasonable request. Proteomics and RNAseq data have been deposited to Zenodo and are available at access number 11383989.

## Supplementary

**Extended Figure 1.**
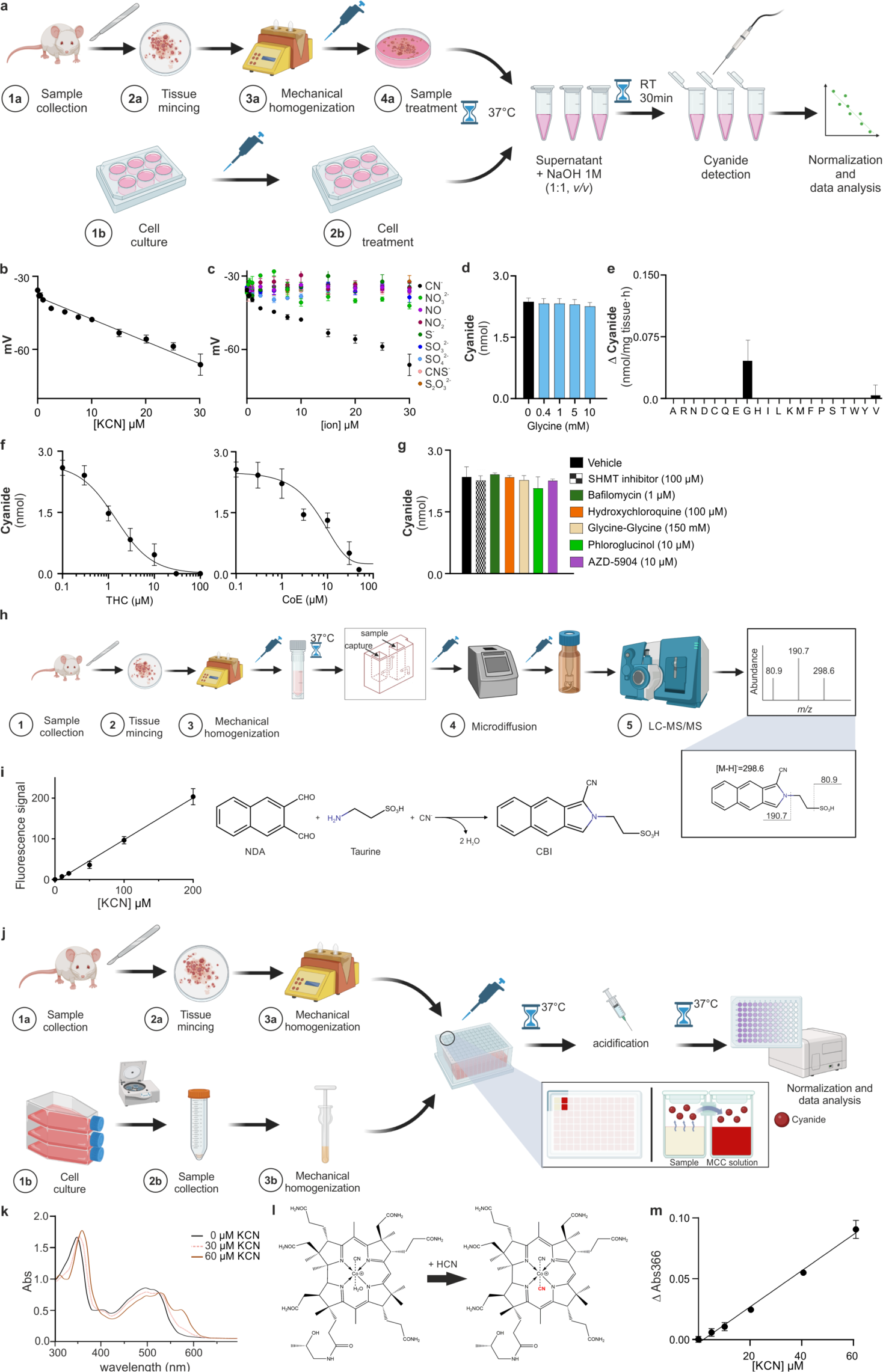
HCN detection methods used. **(a-g)** The electrochemical method **(h-i)** the LC-MS/MS method and **(j-m)** the monocyano-cobinamide (MCC) method. **(a)** Workflow of HCN detection using the electrochemical method (ECh) used in the current study (created with BioRender.com). RT: room temperature. **(b)** Standard curve using authentic cyanide (KCN) prepared in NaOH 0.5 M in closed Eppendorf tubes and measured after incubation for 30 min at room temperature. **(c)** Ion-selectivity of the method. **(d)** Increasing concentrations of glycine do not interfere with KCN detection **(e)** Δ Increase of HCN generation as compared to the vehicle measured in presence of different proteinogenic amino acids (10 mM) in mouse liver tissue. **(f)** Decrease of free HCN levels in presence of 0-100 µM HCN scavengers trihistidyl-cobinamide (THC) or dicobalt edetate (CoE). **(g)** Increasing concentration of different pharmacological modulators – SHMT inhibitor (iSHMT, 100 µM), bafilomycin (1 µM), hydroxychloroquine (100 µM), Glycine-glycine (150 mM), phloroglucinol (10 µM) and 1,2,3,9-tetrahydro-3-[[(2R)-tetrahydro-2-furanyl]methyl]-2-thioxo-6H-purin-6-one (AZD-5904, 10 µM) used in the current study do not interfere with cyanide detection. Data are presented as mean values ± SEM of at least n=5 independent experiments. **(h)** Workflow of HCN detection using LC-MS/MS (created with BioRender.com). **(i)** Standard-curve of authentic cyanide (KCN) and reaction of HCN with 2,3-naphtalenedialdehyde (NDA) and taurine to produce the fluorescent product 1-cyano benzoisoindole (CBI). **(j)** Workflow of HCN detection using the monocyano-cobinamide (MCC) method (created with BioRender.com). **(k)** UV-vis absorption spectra of MCC exposed to HCN evolved from increasing concentrations of KCN. **(l)** Molecular structure of MCC which, in presence of HCN, is converted to dicyano-cobinamide. **(m)** Standard curve of authentic cyanide (KCN) in the MCC method.

**Extended Figure 2.**
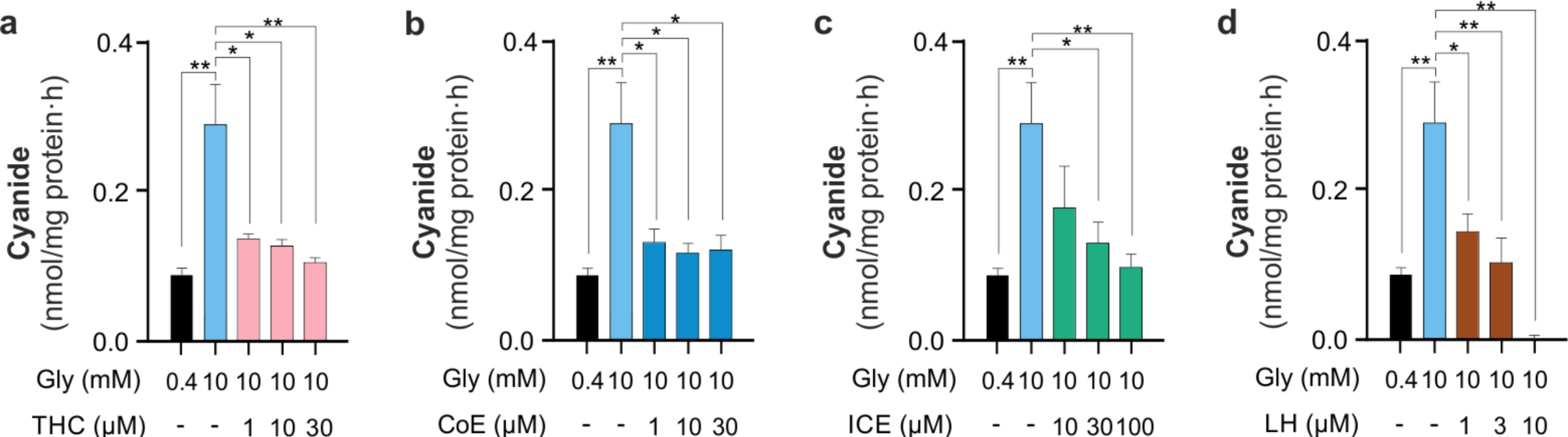
Modulation of HCN levels in HepG2 cells. Increasing concentrations of **(a)** the HCN scavenger trihistidyl-cobinamide (THC, 0-30 µM), **(b)** the HCN scavenger dicobalt edetate (CoE, 0-30 µM) or **(c)** the glycine transporter GlyT-1 inhibitor iclepertin (ICE 10-100 µM) and **(d)** SHMT inhibitor lometrexol hydrate (LH, 1-10 µM) abrogate glycine-mediated stimulation of HCN generation from HepG2 cells, as measured by electrochemical method. Data are presented as mean values ± SEM of at least n=5 independent experiments. *p<0.05 or **p<0.01 indicate significant differences.

**Extended Figure 3.**
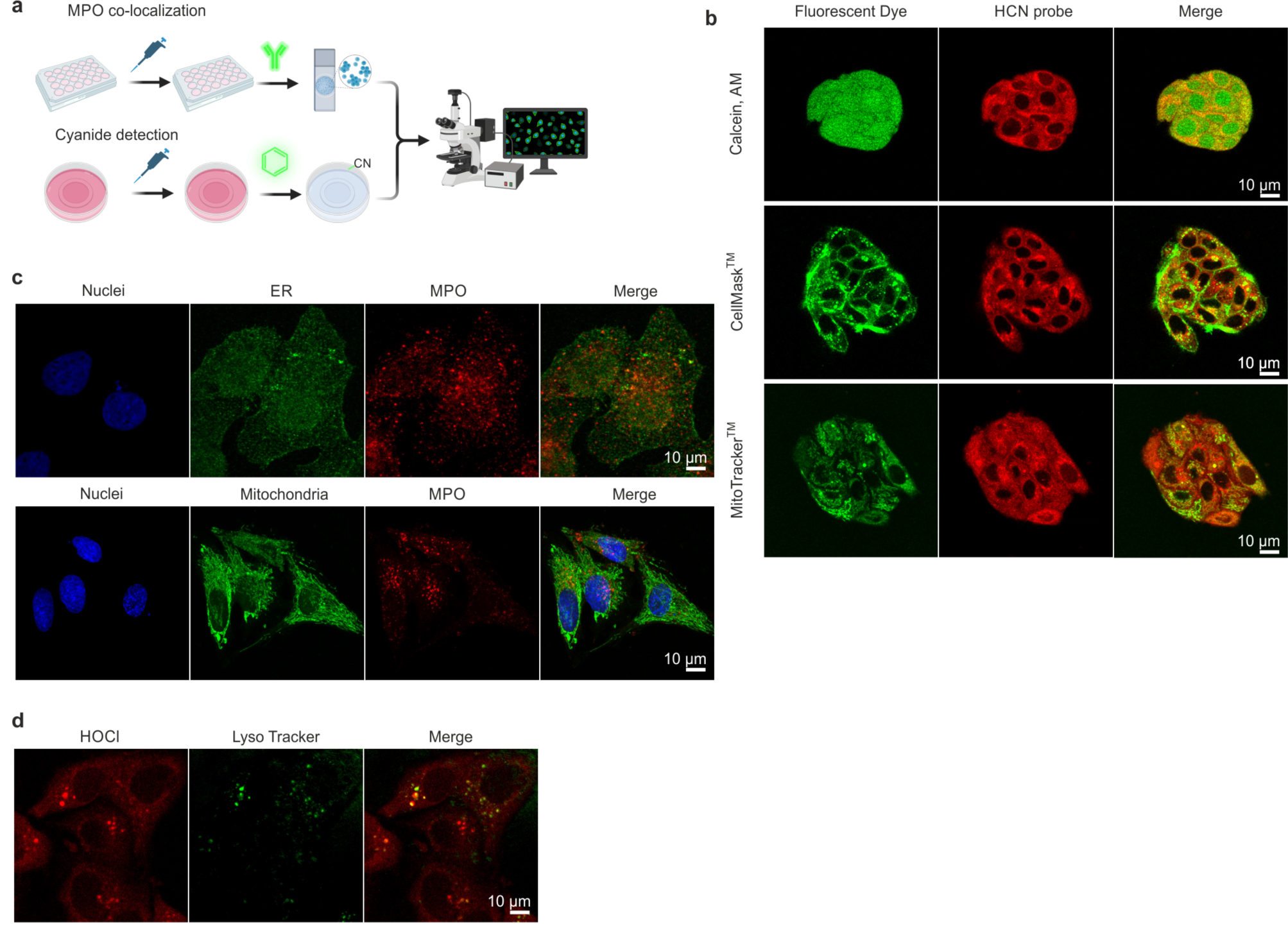
Confocal microscopy analysis of cyanide and HOCl generation in HepG2 cells. **(a)** Workflow of HCN detection using confocal microscopy (created with BioRender.com). MPO: myeloperoxidase. **(b)** Confocal microscopy of HepG2 cells incubated with 10 µM selective HCN probe for 1 h incubation together with other cell-permeant dyes (10 µM Calcein AM, or 1 µM CellMask™ Green Actin Tracking Stain, or 200 nM MitoTracker Deep Red FM) for 30 minutes at 37 °C and 5% CO_2_. At the end of the incubation, cells were washed 3-times and visualized using confocal microscope. Following excitation and emission spectra were used: HCN probe (Ex 405 nm/Em 584-620), Calcein AM (Ex 488/Em 517), CellMask™ Plasma Membrane Stain (Ex 488/Em 535) MitoTrackerDeepRed (Ex 644/Em 665). **(c)** Colocalization of MPO with various subcellular organelles. Representative confocal microscopy image of HepG2 showing that MPO does not localize to the endoplasmic reticulum (ER) nor mitochondria. 500 nM ER tracker green was employed for ER detection. Primary antibody anti-MPO antibody (mouse, dilution 1:10,000, Sigma-Aldrich) followed by incubation with appropriate secondary antibody goat anti-rabbit IgG (H+L) Highly Cross-Adsorbed Secondary Antibody Alexa Fluor Plus 647 (1:1,000 dilution) was used for detection of MPO. 200 nM MitoTracker Deep Red was employed for mitochondria detection. DAPI (Molecular Probes, 5 µg/ml) was used for the detection of cellular nuclei. Images were visualized using Leica 8 STELLARIS Falcon at 63x magnification. For co-localization with ER, ER tracker was visualized at Ex504/Em511 nm and MPO at Ex647/Em 665nm. For co-localization with mitochondria, MitoTracker was visualized at Ex647/Em 665nm and MPO at Ex 568nm. **(d)** Co-localization of HOCl with lysosomes. Representative confocal microscopy image of HepG2 showing that HOCl localizes with lysosomes. HepG2 cells were loaded with 10 µM Chemosensor P in HBSS for 1 h together with 50 nM LysoTrackerGreen for 30 min at 37 °C and 5% CO_2_. At the end of the incubation, cells were washed 3-times and visualized using confocal microscope. Following excitation and emission spectra were used: HOCl probe (Ex 405 nm/Em 450-550) and LysoTrackerGreen Ex488/517.

**Extended Figure 4.**
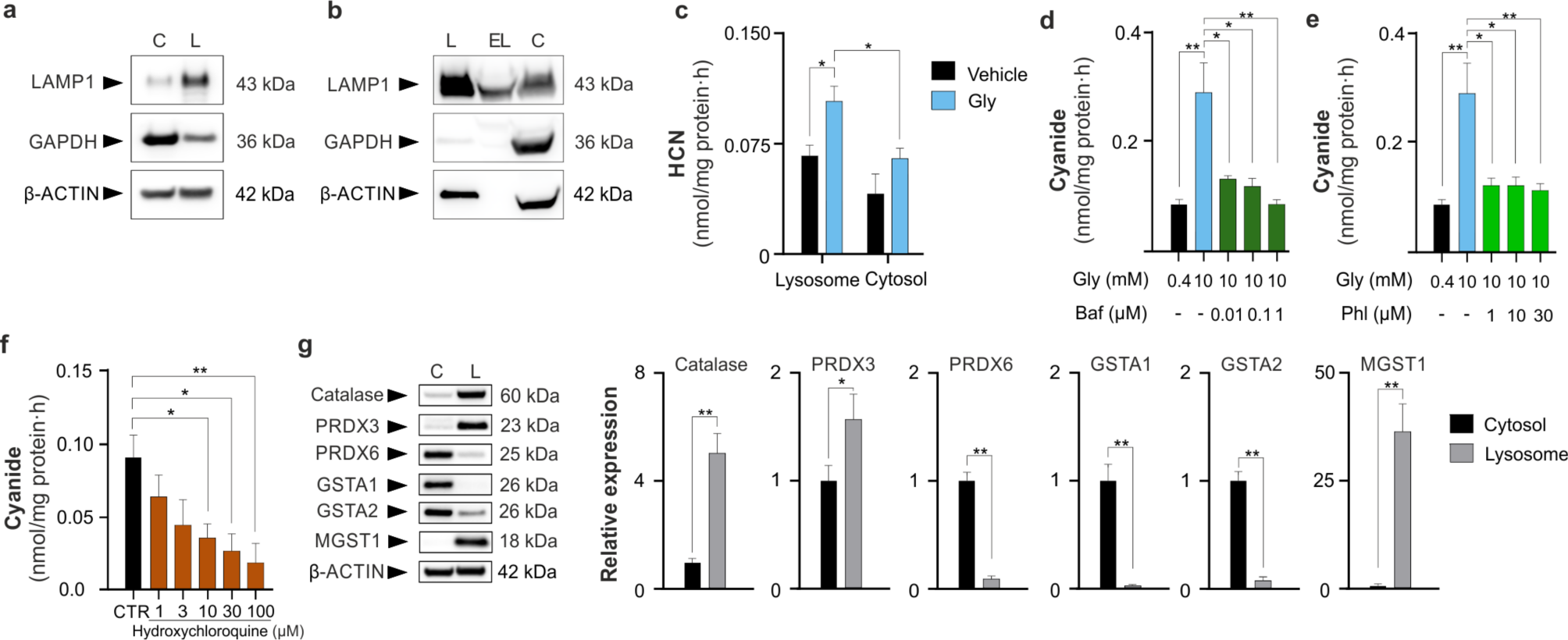
Mechanisms of lysosomal HCN production. **(a)** Western blot of cytosolic (C) *vs* lysosomal (L) fractions from HepG2 cells. LAMP-1: lysosomal associated membrane protein 1; GAPDH: glyceraldehyde 3-phosphate dehydrogenase. **(b)** Western blot of lysosomal (L), extra-lysosomal (EL) and cytosolic (C) fractions from mouse liver. **(c)** HCN detection in the lysosomal *vs* cytosolic fractions of HepG2 cells, with or without incubation with glycine (10 mM), as measured with the LC-MS/MS method. **(d)** Inhibitory effect of the lysosomal proton pump inhibitor bafilomycin (Baf, 0.1- 1 µM) on cyanide production in HepG2 cells in the presence of increasing concentration (10 mM) of glycine (Gly). **(e)** Inhibitory effect of the peroxidase inhibitor phloroglucinol (Phl, 1- 30 µM) on cyanide production in HepG2 cells. **(f)** Inhibitory effect of the lysosomal deacidifier hydroxychloroquine (1-100 µM) on cyanide production in HepG2 cells. **(e)** Western blots of the expression of catalase and various peroxidases – catalase, peroxiredoxin3 (PRDX3), peroxiredoxin6 (PRDX6), glutathione S-transferase alpha1 (GSTA1), glutathione S-transferase alpha2 (GSTA2), and microsomal glutathione S-transferase1 (MGST1) – in cytosolic (C) vs. lysosomal (L) fractions of HepG2 cells. Data are presented as mean values ± SEM of n=5 independent experiments. *p<0.05 and **p<0.01 indicate significant differences.

**Extended Figure 5.**
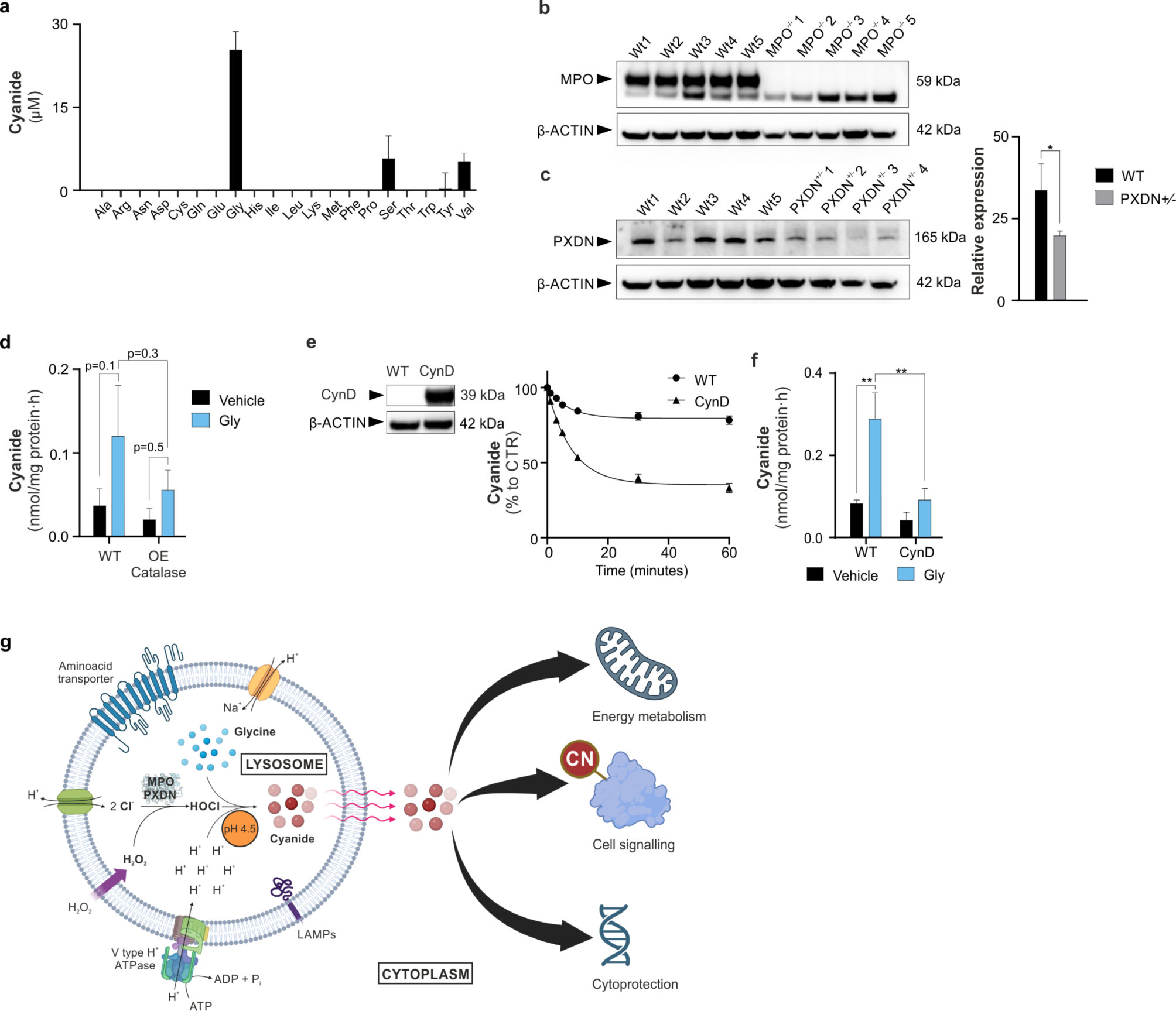
HOCl-catalyzed lysosomal HCN generation. **(a)** HCN detection with electrochemical method using equimolar concentrations of HOCl mixed with different proteinogenic amino acids, at pH 4.5 in 50 mM sodium citrate buffer **(b,c)** myeloperoxidase (MPO) and peroxidasin (PXDN) expression in MPO^-/-^ and PXDN^+/-^ mice compared to wild-type (WT) control: Western blot of MPO and PXDN from MPO^-/-^ and PXDN^+/-^ mouse liver homogenates, respectively. For MPO, no detectable protein band was found in the MPO^-/-^ mice. **(d)** Detection of HCN in HEK293T cells overexpressing catalase (OE catalase). HCN generation has been measured in HEK293T cells overexpressing catalase in the absence or presence of glycine (Gly) supplementation (10 mM) using the ECh method. Data are expressed as presented as mean values ± SEM of at least n=5 independent experiments. *p<0.05 indicates a significant difference for PXDN expression between WT vs. PXDN^+/-^ mice. **(e)** Decomposition of exogenously applied cyanide (KCN) in wild-type HepG2 cells vs. HepG2 cells overexpressing cyanide dihydratase (CynD). **(f)** Cyanide production in wild-type HepG2 cells vs. HepG2 cells overexpressing CynD with or without the addition of Gly (10 mM). **(g)** Cyanide generation in mammalian cells: mechanisms and consequences. Cyanide generation occurs mainly in lysosomes: here, myeloperoxidase (MPO) or peroxidasin (PXDN) catalyzes the production of HOCl from hydrogen peroxide (H_2_O_2_) and chloride (Cl^-^). At physiological lysosomal pH (4.5), glycine is chlorinated by HOCl to generate N,N-dichloroglycine, the decomposition of which yields cyanide, CO_2_ and Cl. Cyanide diffuses through the lysosomal membrane to the cytosol where it acts as a signaling-molecule (in part through S-cyanylation of target proteins), stimulates bioenergetics and provides cytoprotective effects.

**Extended Figure 6.**
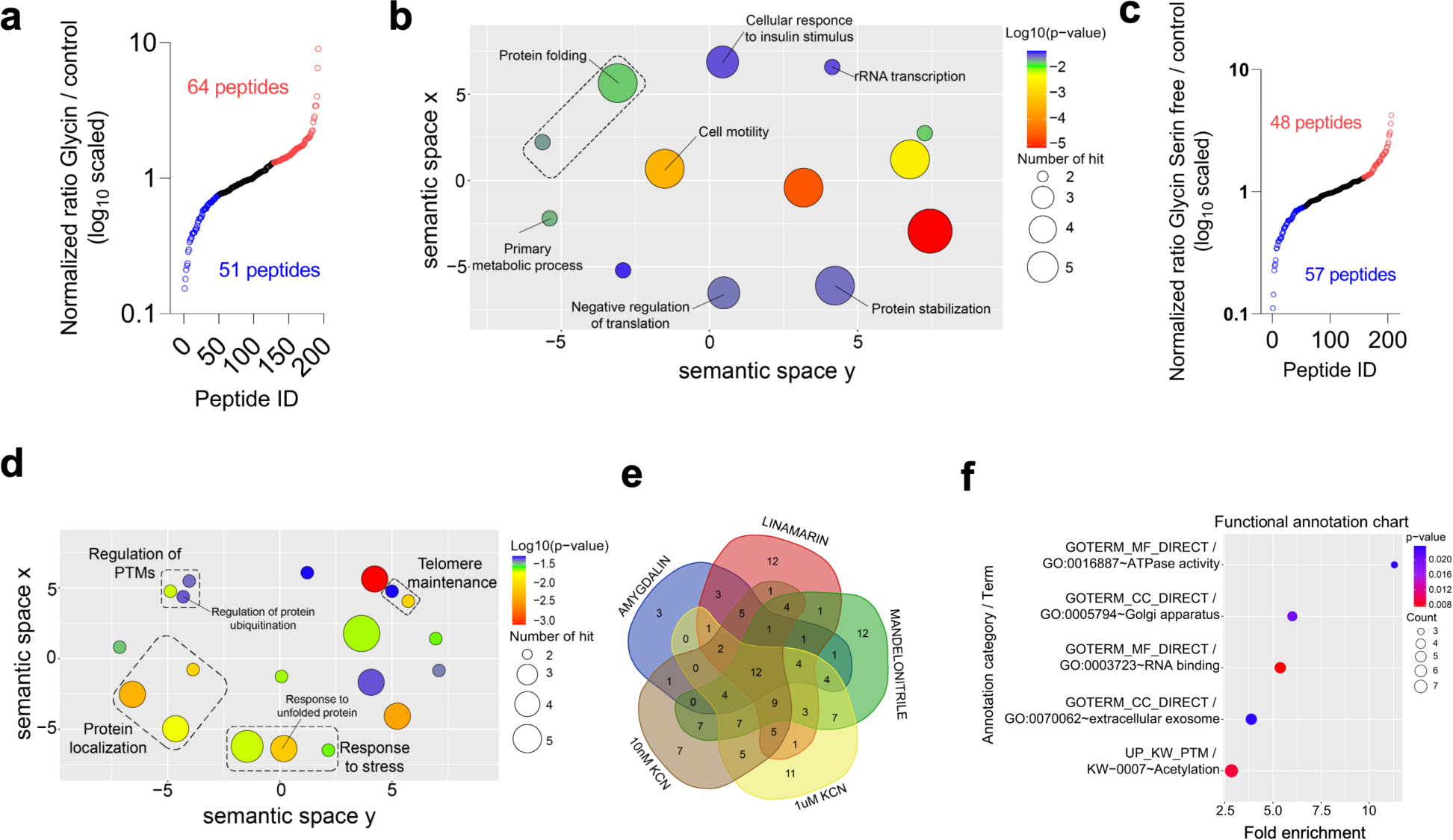
S-cyanylome remodeling caused by glycine or various cyanide donors. **(a)** The proteome-wide and site-specific changes in *S*-cyanylation in HepG2 cells supplemented with 10 mM glycine. **(b)** GO term enrichment (biological process) of the proteins whose S-cyanylation is significantly increased in HepG2 cells treated with 10 mM glycine. Benjamini adjusted p-value threshold is 0.01. Circle dimensions denote the protein count within specific GO terms, while color gradients communicate the degree of significance. **(c)** The proteome-wide and site-specific changes in *S*-cyanylation in HepG2 cells cultured in Ser/Gly-free medium. **(d)** GO term enrichment (biological process) of the proteins whose S-cyanylation significantly decreased in HepG2 cells grown in Ser/Gly-free medium. Benjamini adjusted p-value threshold is 0.01. Circle dimensions denote the protein count within specific GO terms, while color gradients communicate the degree of significance. PTM: posttranslational modification. **(e)** Venn diagram of proteins whose S-cyanylation was found to be significantly increased in HepG2 cells treated with 1 nM KCN, 10 nM KCN, 100 µM linamarin, 30 µM amygdalin or 100 µM mandelonitrile. **(f)** GO term enrichment analysis of 12 proteins found to be commonly increasing in response to all 5 treatments.

**Extended Figure 7.**
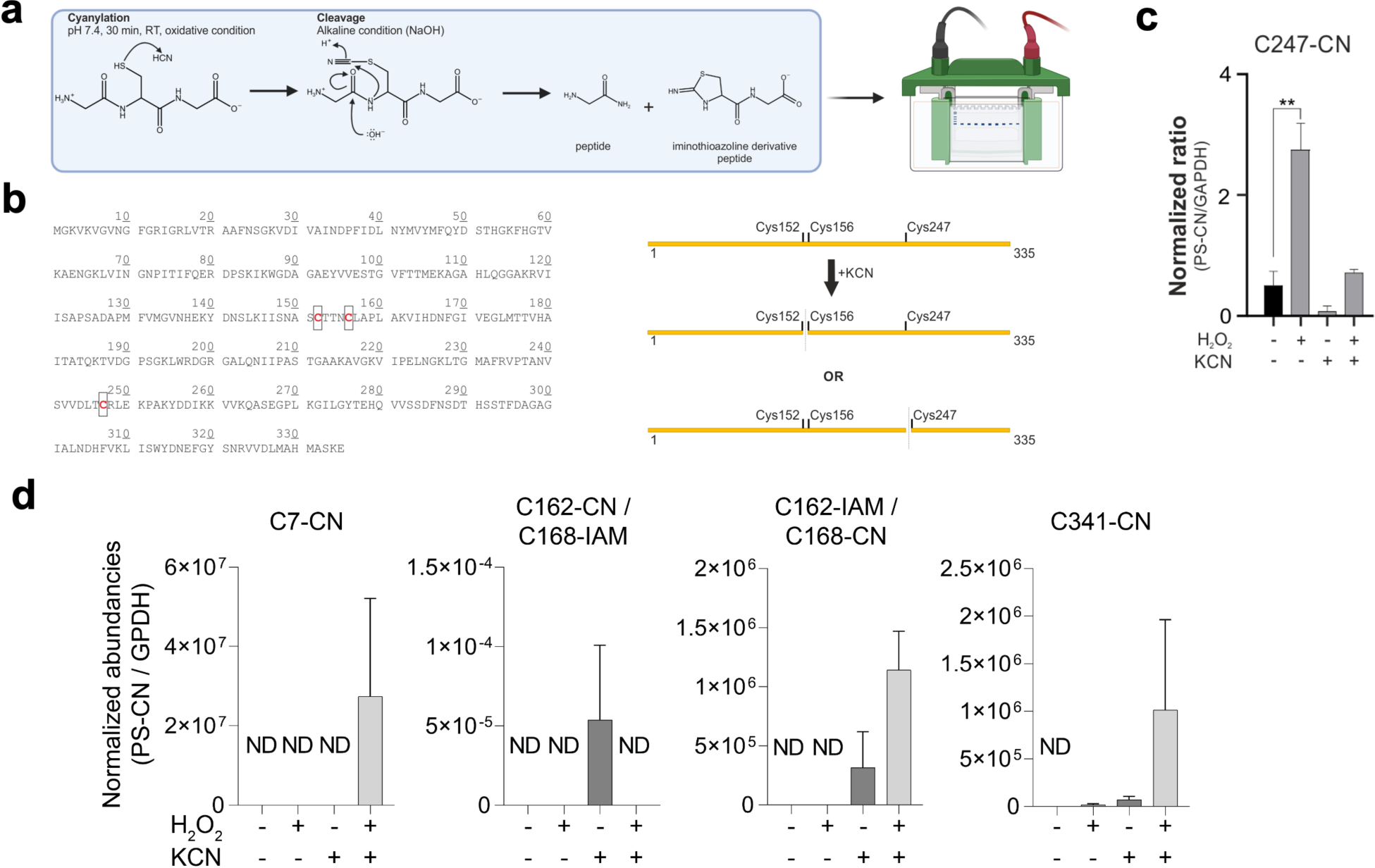
S-Cyanylation of glyceraldehyde 3-phosphate dehydrogenase (GAPDH) and Glycerol-3-phosphate dehydrogenase (GPDH). **(a)** Chemical cleavage of polypeptide’s backbone occurs after S-cyanylation of target cysteine residues under alkaline conditions. Cleavage is then detected by SDS-PAGE followed by Coomassie-staining. **(b)** Human GAPDH contains three cysteine residues. S-cyanylation of GAPDH results in the generation of two visible bands that are consistent with one single cleavage reaction and one S-cyanylation site. **(c)** MS label-free quantification of S-cyanylation of C247 peptide. Peptide intensity is normalized to the total GAPDH in each sample. 0.55 µM GAPDH was treated with 10 µM KCN 10 µM H_2_O_2_ or the combination of KCN and H_2_O_2_. **(d)** MS label-free quantification of GPDH treated with 10 µM KCN 10 µM H_2_O_2_ or the combination of KCN and H_2_O_2_. ND: not detectable. Data represent mean values ± SEM of at least n=3 independent experiments; *p<0.05 or **p<0.01 indicate significant differences.

**Extended Figure 8.**
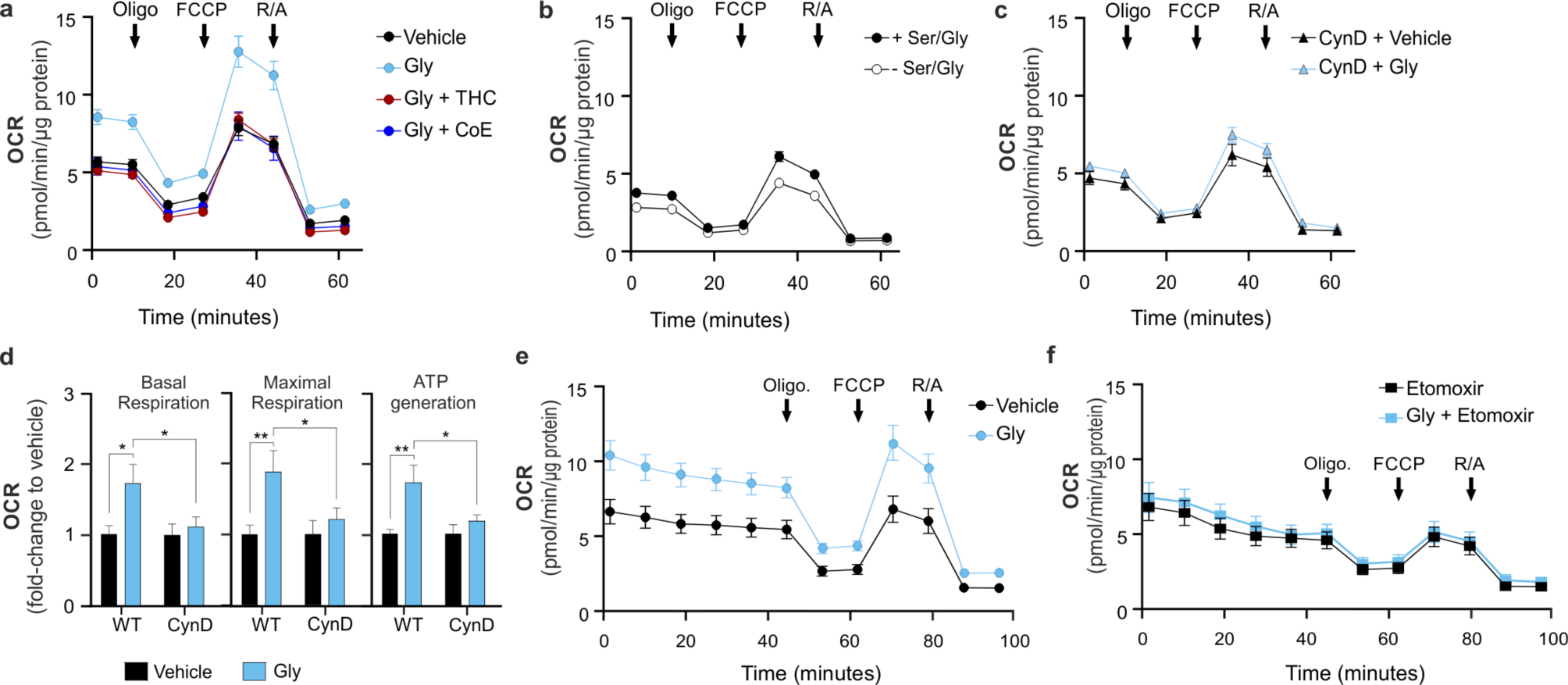
Endogenous HCN modulates cellular bioenergetics in HepG2 cells: extracellular flux analysis. **(a)** Glycine-dependent stimulation of cellular bioenergetics is abrogated by HCN scavengers. Extracellular flux analysis of HepG2 cells incubated with 10 mM glycine (Gly) for 24 h followed by 3 h with 10 µM of the cyanide scavengers (trihistidyl-cobinamide (THC) or dicobalt edetate (CoE). OCR: oxygen consumption rate. **(b)** HepG2 cells incubated with serine/glycine free medium (-Ser/Gly) or with serine/glycine free medium re-supplemented with 0.4 mM serine and 0.4 mM glycine (+Ser/Gly). **(c,d)**. Glycine-dependent stimulation of cellular bioenergetics is abrogated by overexpression of the HCN-catabolizing enzyme cyanide dihydratase (CynD) in HepG2 cells, compared to normal control wild-type (WT) cells. Extracellular flux analysis. **(e,f)** Free fatty acid oxidation extracellular flux analysis of (**e**) control HepG2 cells and (**f**) HepG2 cells pretreated with 4 µM etomoxir (carnitine palmitoyl transferase-1 inhibitor). In both cases, cells were pre-incubated in presence of 10 mM glycine or vehicle for 24 h. Arrows in panels **a**, **b**, **c**, **e** and f represent the times of the addition of the ATP synthase inhibitor oligomycin, the uncoupling agent carbonyl cyanide-4-(trifluoromethoxy)phenylhydrazone (FCCP) and the combined addition of the mitochondrial Complex I inhibitor rotenone and the mitochondrial Complex III inhibitor antimycin (R/A) in the Extracellular Flux Analysis protocol. Data represent original plot normalized with total protein of n=5 independent experiments ± SEM. *p<0.05 indicates significant differences.

**Extended Figure 9.**
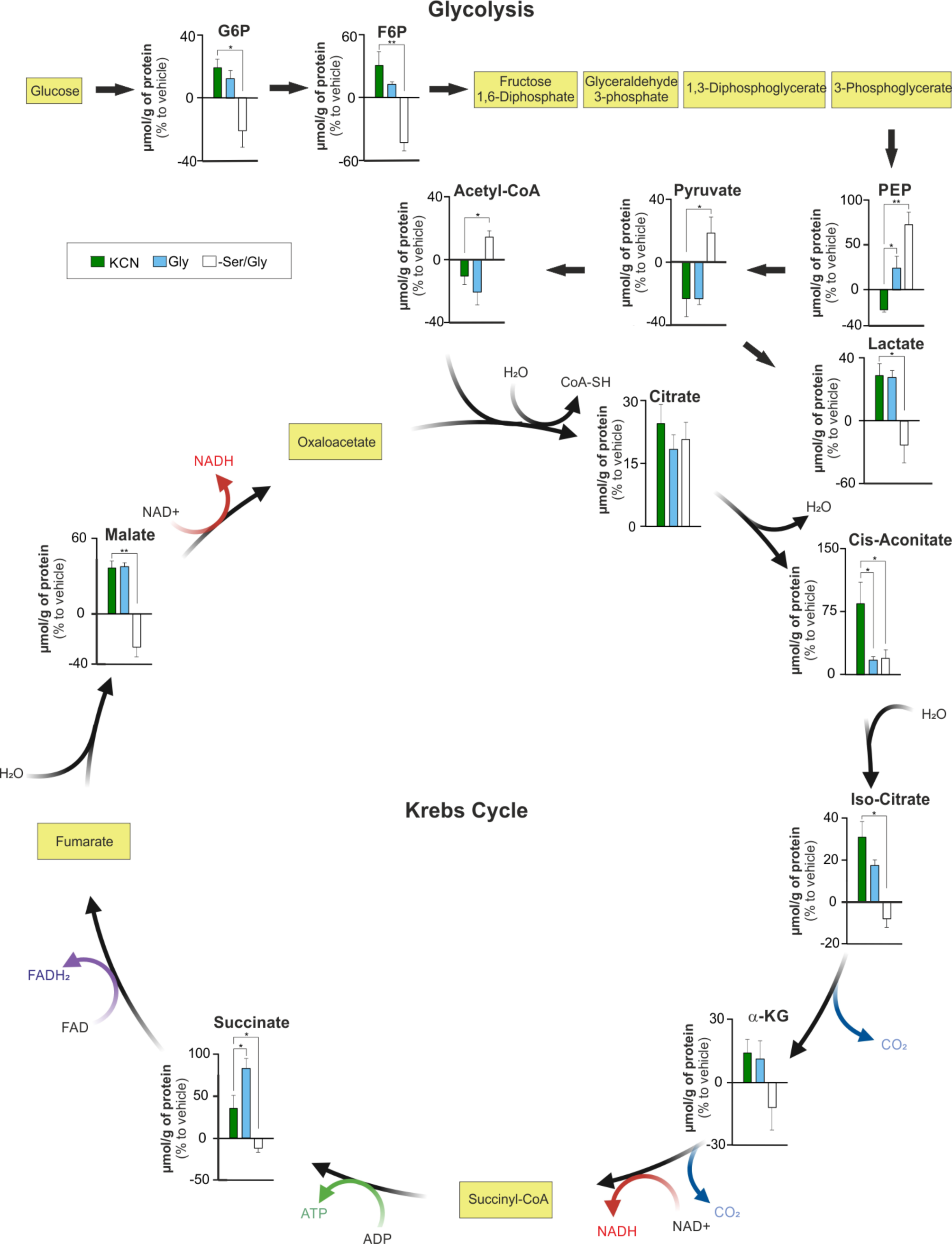
Effect of KCN, glycine supplementation or serine/glycine deprivation on glycolysis and TCA cycle. Metabolite levels analyzed by LCMS in HepG2 cells cultured with media supplemented with 10 nM KCN, 10 mM glycine (Gly) or under deprivation of serine and glycine (-Ser/Gly). Metabolites in yellow squares were not detected by the method used. Data show mean of n=3 independent experiments ±SEM; *p<0.05 or **p<0.01 indicate significant differences.

**Extended Figure 10.**
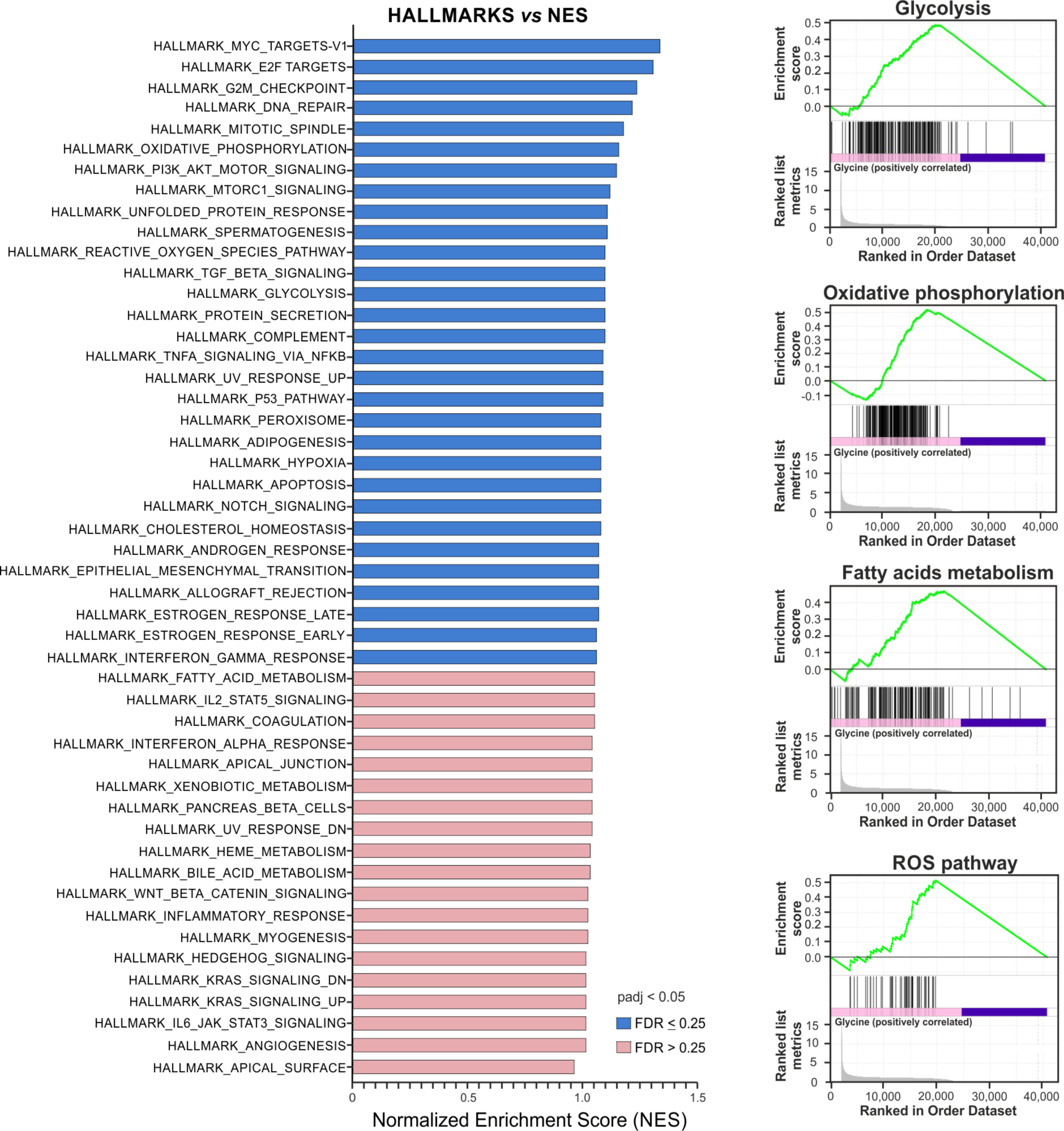
Glycine-induced transcriptomic changes. Gene Set Enrichment Analysis (GSEA), using the hallmark pathway gene sets of HepG2 cells incubated with 10 mM glycine for 24 h compared to vehicle. Data were obtained from RNAseq of n=3 independent replicates per group.

**Extended Data Table S1.** Cyanylated and carbamidomethylated proteins in mouse liver tissues. Liver tissue was treated with vehicle of 10 mM glycine.

**Extended Data Table S2.** Protein cyanylation in mouse liver tissues. Liver tissue was treated with vehicle or 10 mM ’heavy’ glycine.

**Extended Data Table S3.** Protein cyanylation in HepG2 cells. HepG2 cells were grown in the medium supplemented with 10 mM glycine or grown in serine/glycine-free medium.

**Extended Data Table S4.** Protein cyanylation in HepG2 cells. HepG2 cells were treated with various cyanide releasers (amygdalin, linamarin, mandelonitrile) or with KCN.

**Extended Data Table S5.** Up-regulated genes of the glycolytic pathway detected by RNAseq analysis in HepG2 cells.

**Extended Data Table S6.** Up-regulated genes of the oxidative phosphorylation pathway detected by RNAseq analysis in HepG2 cells.

**Extended Data Table S7.** Up-regulated genes of the fatty acid metabolism detected by RNAseq analysis in HepG2 cells.

**Extended Data Table S8.** Up-regulated genes of the reactive oxygen species pathway detected by RNAseq analysis in HepG2 cells.

**Extended Data Table S9.** Up- and down regulated genes detected by RNAseq analysis after TST silencing in HepG2 cells.

**Extended Data Table S10.** Up- and down regulated genes detected by RNAseq analysis after TST overexpression in HepG2 cells.

**Extended Data Table S11.** Comparison of the biological production and cellular action of the gasotransmitters nitric oxide (NO), carbon monoxide (CO), hydrogen sulfide (H2S) and hydrogen cyanide (HCN).

## Footnotes

* Cyanide is a weak acid (pKa = 9.2). At physiological pH, approximately 95% of this species exists in the volatile undissociated form (HCN) and 5% in the dissociated form (cyanide, CN^-^). In this article, we refer to all species in biological systems as “cyanide.” As described in Methods, we measured cyanide by complementary methods, either after alkalinization to generate CN^-^ (electrode method) or acidification to generate HCN (Cyanalyzer and spectrophotometric method).

b Hydroxychloroquine and chloroquine are lipophilic and lysosomotropic drugs, that penetrate cell membranes, and accumulate in the acidic lysosomes. These alkaline compounds increase the pH in lysosomes – from the normal values of 4.7–4.8 to 6.

## References

1. Wang, R. Gasotransmitters: growing pains and joys. Trends Biochem. Sci. 39, 227–232 (2014).

2. Kelm, M. Nitric oxide metabolism and breakdown. Biochim. Biophys. Acta 1411, 273–289 (1999).

3. Moncada, S. Palmer, R.M., & Higgs, E.A. Nitric oxide: physiology, pathophysiology, and pharmacology. Pharmacol Rev. 43, 109–42 (1991).

4. Wang, R. Physiological implications of hydrogen sulfide: a whiff exploration that blossomed. Physiol. Rev. 92, 791–896 (2012).

5. Wu, L. & Wang, R. Carbon monoxide: endogenous production, physiological functions, and pharmacological applications. Pharmacol. Rev. 57, 585–630 (2005).

6. Cirino, G., Szabo, C. & Papapetropoulos, A. Physiological roles of hydrogen sulfide in mammalian cells, tissues, and organs. Physiol. Rev. 103, 31–276 (2023).

7. Pacher, P., Beckman, J.S. & Liaudet, L. Nitric oxide and peroxynitrite in health and disease. Physiol. Rev. 87, 315–424 (2007).

8. Hendry-Hofer, T.B., et al., A review on ingested cyanide: risks, clinical presentation, diagnostics, and treatment challenges. J. Med. Toxicol. 15, 128–133 (2019).

9. McAnalley, B.H., Lowry, W.T., Oliver, R.D., Garriott, J.C. Determination of inorganic sulfide and cyanide in blood using specific ion electrodes: application to the investigation of hydrogen sulfide and cyanide poisoning. J. Anal. Toxicol. 3, 111–114 (1979).

10. Alluhayb, A.H., Severance, C., Hendry-Hofer, T., Bebarta, V.S., Logue, B.A. Concurrent determination of cyanide and thiocyanate in human and swine antemortem and postmortem blood by high-performance liquid chromatography-tandem mass spectrometry. Anal. Bioanal. Chem. 415, 6595–6609. (2023).

11. Blackledge, W.C. et al., New facile method to measure cyanide in blood. Anal. Chem. 82, 4216–4221 (2010).

12. Broderick, K.E. et al., Cyanide detoxification by the cobalamin precursor cobinamide. Exp. Biol. Med. 231, 641–649 (2006).

13. Marrs, T.C. & Thompson, J.P. The efficacy and adverse effects of dicobalt edetate in cyanide poisoning. Clin. Toxicol. 54, 609–614 (2016).

14. Malkondu, S. & Erdemir, S. Cyanobiphenyl-spiropyrane and -hemicyanine conjugates for cyanide detection in organic/aqueous media through reverse ICT direction: Their practical applications Talanta. 231, 22385 (2021).

15. Malkondu, S., Erdemir, S. & Karakurt, S. Red and blue emitting fluorescent probe for cyanide and hypochlorite ions: Biological sensing and environmental analysis. Dyes Pigm. 174, 108019 (2020).

16. Ducker, G.S. & Rabinowitz, J.D. One-carbon metabolism in health and disease. Cell. Metab. 25, 27–42 (2017).

17. Yoshimori, T., Yamamoto, A., Moriyama, Y., Futai, M. & Tashiro, Y. Bafilomycin A1, a specific inhibitor of vacuolar-type H^+^-ATPase, inhibits acidification and protein degradation in lysosomes of cultured cells. J. Biol. Chem. 266, 17707–17712 (1991)

18. Poole, B. & Ohkuma, S. Effect of weak bases on the intralysosomal pH in mouse peritoneal macrophages. J. Cell. Biol. 90, 665–669 (1981).

19. Frølund, S., Langthaler, L., Kall, M.A. Holm, R. & Nielsen, C.U. Intestinal drug transport via the proton-coupled amino acid transporter PAT1 (SLC36A1) is inhibited by Gly-X(aa) dipeptides. Mol. Pharm. 9, 2761–2769 (2012).

20. Colon, S. et al., Peroxidasin and eosinophil peroxidase, but not myeloperoxidase, contribute to renal fibrosis in the murine unilateral ureteral obstruction model. Am. J. Physiol. Renal Physiol. 316, F360–F371 (2019).

21. Khan, A.A., Rahmani, A.H., Aldebasi, Y.H. & Aly, S.M. Biochemical and pathological studies on peroxidases - an updated review. Glob. J. Health Sci. 6, 87–98 (2014).

22. Jiang, J. et al., *In vivo* bioimaging and detection of endogenous hypochlorous acid in lysosome using a near-infrared fluorescent probe. Anal. Methods 15, 3188–3195 (2023).

23. Tidén, A.K. et al., 2-thioxanthines are mechanism-based inactivators of myeloperoxidase that block oxidative stress during inflammation. J. Biol. Chem. 286, 37578–37589 (2011).

24. Buonvino, S., Arciero, I. & Melino, S. Thiosulfate-cyanide sulfurtransferase a mitochondrial essential enzyme: from cell metabolism to the biotechnological applications. Int. J. Mol. Sci. 23, 8452 (2022).

25. Martínková, L., Veselá, A.B., Rinágelová, A. & Chmátal, M. Cyanide hydratases and cyanide dihydratases: emerging tools in the biodegradation and biodetection of cyanide. Appl. Microbiol. Biotechnol. 99, 8875–8882 (2015).

26. Zgliczyński, J.M. & Stelmaszyńska, T. Hydrogen cyanide and cyanogen chloride formation by the myeloperoxidase-H_2_O_2_-Cl-system. Biochim. Biophys. Acta 567, 309–314 (1979).

27. García, I., Arenas-Alfonseca, L., Moreno, I., Gotor, C. & Romero, L.C. HCN regulates cellular processes through posttranslational modification of proteins by S-cyanylation. Plant Physiol. 179, 107–123 (2019).

28. Fasco, M.J. et al., Cyanide adducts with human plasma proteins: albumin as a potential exposure surrogate. Chem. Res. Toxicol. 20, 677–684 (2007).

29. Youso, S.L., Rockwood, G.A., Lee, J.P. & Logue, B.A. Determination of cyanide exposure by gas chromatography-mass spectrometry analysis of cyanide-exposed plasma proteins. Anal. Chim. Acta 677, 24–28 (2010).

30. Randi, E.B., Zuhra, K., Pecze, L., Panagaki, T. & Szabo, C. Physiological concentrations of cyanide stimulate mitochondrial Complex IV and enhance cellular bioenergetics. Proc. Natl. Acad. Sci. USA. 118, e2026245118 (2021).

31. Vinnakota, C.V. et al., Comparison of cyanide exposure markers in the biofluids of smokers and non-smokers. Biomarkers. 17, 625–33 (2012).

32. Nowak, M., Chuchra, P. & Paprocka, J. Nonketotic hyperglycinemia: insight into current therapies. J. Clin. Med. 11, 3027 (2022).

33. Correia, S.C. et al., Cyanide preconditioning protects brain endothelial and NT2 neuron-like cells against glucotoxicity: role of mitochondrial reactive oxygen species and HIF-1α. Neurobiol. Dis. 45, 206–218 (2012).

34. Zuhra, K. & Szabo, C. The two faces of cyanide: an environmental toxin and a potential novel mammalian gasotransmitter. FEBS J. 289, 2481–2515 (2022).

35. Díaz-Rueda, P., Morales de Los Ríos, L., Romero, L.C. & García, I. Old poisons, new signaling molecules: the case of HCN. J. Exp. Bot. erad317 (2023).

36. Bhandari, R.K. et al., Simultaneous determination of cyanide and thiocyanate in plasma by chemical ionization gas chromatography mass-spectrometry (CI-GC-MS). Anal. Bioanal. Chem. 404, 2287–2294 (2012).

37. Houten, S.M., Violante, S., Ventura, F.V. & Wanders, R.J. The biochemistry and physiology of mitochondrial fatty acid β-oxidation and its genetic disorders. Annu. Rev. Physiol. 78, 23–44 (2016).

38. Alves, A., Bassot, A., Bulteau, A.L., Pirola, L. & Morio, B. Glycine metabolism and its alterations in obesity and metabolic diseases. Nutrients 11, 1356 (2019).

39. Jois, M., Hall, B., Fewer, K. & Brosnan, J.T. Regulation of hepatic glycine catabolism by glucagon. J. Biol. Chem. 264, 3347–51 (1989).

40. Abu-Remaileh, M. et al., Lysosomal metabolomics reveals V-ATPase- and mTOR-dependent regulation of amino acid efflux from lysosomes. Science 358, 807–813 (2017).

41. Petrat, F., Boengler, K., Schulz, R. & de Groot, H. Glycine, a simple physiological compound protecting by yet puzzling mechanism(s) against ischaemia-reperfusion injury: current knowledge. Br. J. Pharmacol. 165, 2059–2072 (2012).

42. Milazzo, S. & Horneber, M. Laetrile treatment for cancer. Cochrane Database Syst. Rev. 2015, CD005476 (2015).

43. Duranski, M.R. et al., Cytoprotective effects of nitrite during in vivo ischemia-reperfusion of the heart and liver. J. Clin. Invest. 115, 1232–40 (2005).

44. Elrod, J.W. et al., Hydrogen sulfide attenuates myocardial ischemia-reperfusion injury by preservation of mitochondrial function. Proc. Natl. Acad. Sci. USA. 104, 15560–5 (2007).

45. Blumer, C. & Haas, D. Mechanism, regulation, and ecological role of bacterial cyanide biosynthesis. Arch. Microbiol. 173, 170–177 (2000)

46. Boter, M. & Diaz, I. Cyanogenesis, a plant defence strategy against herbivores. Int. J Mol Sci. 24, 6982 (2023).

47. Olson, K.R. & Straub, K.D. The role of hydrogen sulfide in evolution and the evolution of hydrogen sulfide in metabolism and signaling. Physiology (Bethesda). 31, 60–72 (2016).

48. Das, T., Ghule, S. & Vanka, K. Insights into the origin of life: Did it begin from HCN and H_2_O? ACS Cent. Sci. 5, 1532–1540 (2019).

49. Zhu, J., Ligi, S. & Yang, G. An evolutionary perspective on the interplays between hydrogen sulfide and oxygen in cellular functions. Arch. Biochem. Biophys. 707, 108920 (2021).

50. Yadav, M., Pulletikurti, S., Yerabolu, J.R. & Krishnamurthy, R. Cyanide as a primordial reductant enables a protometabolic reductive glyoxylate pathway. Nat. Chem. 14, 170–178 (2022).

51. Cortese-Krott, M.M., et al., The reactive species interactome: evolutionary emergence, biological significance, and opportunities for redox metabolomics and personalized medicine. Antioxid Redox Signal. 27, 684–712 (2017).

52. Vignane, T. & Filipovic, M.R. Emerging chemical biology of protein persulfidation. Antioxid Redox Signal. 39, 19–39 (2023).

53. Szabo, C. Gaseotransmitters: new frontiers for translational science. Sci. Transl. Med. 2, 59ps54 (2010).

54. Zivanovic, J., et al., Selective persulfide detection reveals evolutionarily conserved antiaging effects of S-sulfhydration. Cell Metab. 31, 207 (2020).

55. Beck, K.F. & Pfeilschifter, J. Gasotransmitter synthesis and signalling in the renal glomerulus. Implications for glomerular diseases. Cell Signal. 77, 109823 (2021).

56. Bieza, S., Mazzeo, A., Pellegrino, J. & Doctorovich, F. H_2_S/thiols, NO•, and NO^-^/HNO: interactions with iron porphyrins. ACS Omega 7, 1602–1611 (2022).

57. Bibli S-I. Tyrosine phosphorylation of eNOS regulates myocardial survival after an ischaemic insult: role of PYK2. Cardiovasc. Res. 113, 926–937 (2017).

58. Kelestemur, T., Németh, Z.H., Pacher, P., Antonioli, L. & Haskó, G. A_2A_ adenosine receptors regulate multiple organ failure after hemorrhagic schock in mice. Shock 58, 321–331 (2022).

59. Bains, H. & Singh, R. Isolation of autophagic fractions from mouse liver for biochemical analyses. STAR Protoc. 2, 100730 (2021).

60. Beermann, C., Wunderli-Allenspach, H., Groscurth, P. & Filgueira, L. Lipoproteins from Borrelia burgdorferi applied in liposomes and presented by dendritic cells induce CD8(+) T-lymphocytes in vitro. Cell. Immunol. 201, 124–31 (2000).

61. Zlosnik, J.E. & Williams, H.D. Methods for assaying cyanide in bacterial culture supernatant. Lett. Appl. Microbiol. 38, 360–5 (2004).

62. Nair, C.G., Ryall, B. & Williams, H.D. Cyanide measurements in bacterial culture and sputum. Methods Mol. Biol. 1149, 325–36 (2014).

63. Eiserich, J.P., et al., Quantitative assessment of cyanide in cystic fibrosis sputum and its oxidative catabolism by hypochlorous acid. Free Radic. Biol. Med. 129, 146–154 (2018).

64. Bortey-Sam, N., et al., Diagnosis of cyanide poisoning using an automated, field-portable sensor for rapid analysis of blood cyanide concentrations. Anal. Chim. Acta 1098, 125–132 (2020).

65. Alluhayb, A.H., Severance, C., Hendry-Hofer, T., Bebarta, V.S. & Logue, B.A. Can the cyanide metabolite, 2-aminothiazoline-4-carboxylic acid, be used for forensic verification of cyanide poisoning? Forensic Toxicol. in press, (2024)

66. Greenawald, L.A., Boss, G.R., Snyder, J.L., Reeder, A. & Bell, S. Development of an inexpensive RGB color sensor for the detection of hydrogen cyanide gas. ACS Sens. 2, 1458–1466 (2017)

67. Kosower, N.S. & Kosower, E.M. Diamide: an oxidant probe for thiols, Methods Enzymol. 251, 123–33 (1995).

68. Fasco, M.J., et al., Cyanide adducts with human plasma proteins: albumin as a potential exposure surrogate. Chem. Res. Toxicol. 20, 677–84 (2007).

69. Yang, H., Liu, N. & Liu, S. Determination of peptide and protein disulfide linkages by MALDI mass spectrometry, Top Curr. Chem. 331, 79–116 (2013).

70. Zuhra, K., et al., Mechanism of cystathionine-β-synthase inhibition by disulfiram: The role of bis(N,N-diethyldithiocarbamate)-copper(II). Biochem. Pharmacol. 182, 114267 (2020).

71. Augsburger, F., et al., Role of 3-mercaptopyruvate sulfurtransferase in the regulation of proliferation, migration, and bioenergetics in murine colon cancer cells. Biomolecules 10, 447 (2020).

72. Subramanian, A., et al., Gene set enrichment analysis: a knowledge-based approach for interpreting genome-wide expression profiles. Proc. Natl. Acad. Sci. USA. 102, 15545–50 (2005).

